# Efficacy of CDK4/6 inhibitors in preclinical models of malignant pleural mesothelioma

**DOI:** 10.1101/2021.02.08.430196

**Authors:** Elisabet Aliagas, Ania Alay, Maria Martínez-Iniesta, Miguel Hernández-Madrigal, David Cordero, Mireia Gausachs, Eva Pros, Maria Saigí, Sara Busacca, Annabel J. Sharkley, Alan Dawson, Ramón Palmero, Susana Padrones, Samantha Aso, Ignacio Escobar, Ricard Ramos, Roger Llatjós, August Vidal, Mar Varela, Montse Sánchez-Céspedes, Dean Fennell, Cristina Muñoz-Pinedo, Alberto Villanueva, Xavi Solé, Ernest Nadal

**Affiliations:** Clinical Research in Solid Tumors (CReST) group. Oncobell Program. Bellvitge Biomedical Research Institute (IDIBELL), L’Hospitalet de Llobregat (Barcelona), Spain; Unit of Bioinformatics for Precision Oncology, Catalan Institute of Oncology (ICO), L’Hospitalet de Llobregat (Barcelona), Spain; Molecular Mechanisms and Experimental Therapy in Oncology. Oncobell Program. Bellvitge Biomedical Research Institute (IDIBELL), L’Hospitalet de Llobregat (Barcelona), Spain; Chemoresistance group. Oncobell Program, Bellvitge Biomedical Research Institute (IDIBELL), L’Hospitalet de Llobregat (Barcelona), Spain; Cell Death and Metabolism Group, Oncobell Program, Bellvitge Biomedical Research Institute (IDIBELL), L’Hospitalet de Llobregat (Barcelona), Spain; Consortium for Biomedical Research in Epidemiology and Public Health (CIBERESP), Barcelona, Spain; Cancer Genetics Group, Josep Carreras Leukaemia Research Institute (IJC), Badalona, Barcelona, Spain; Department of Genetics and Genome Biology, Leicester Cancer Research Centre, University of Leicester, Leicester, UK; University of Sheffield Teaching Hospitals, Sheffield, UK; Department of Thoracic Surgery, Glenfield Hospital, Leicester, UK; Department of Medical Oncology, Catalan Institute of Oncology, L’Hospitalet de Llobregat (Barcelona), Spain; Department of Respiratory Medicine, Hospital Universitari de Bellvitge, L’Hospitalet de Llobregat (Barcelona), Spain; Department of Thoracic Surgery, Hospital Universitari de Bellvitge, L’Hospitalet de Llobregat (Barcelona), Spain; Department of Pathology, Hospital Universitari de Bellvitge, L’Hospitalet de Llobregat (Barcelona), Spain; Mesothelioma Research Programme, Department of Genetics and Genome Biology, University of Leicester, Leicester, UK; Department of Basic Nursing, School of Medicine and Health Sciences, Universitat de Barcelona, Campus Bellvitge, L’Hospitalet del Llobregat (Barcelona), Spain; Department of Clinical Sciences, School of Medicine and Health Sciences, Universitat de Barcelona, Campus Bellvitge, L’Hospitalet del Llobregat (Barcelona), Spain

**Keywords:** malignant pleural mesothelioma, CDK4/6 inhibitors, drug therapy, MPM *in vivo* models

## Abstract

There is no effective therapy for patients with malignant pleural mesothelioma (MPM) who progressed to platinum-based chemotherapy and immunotherapy. Here, we investigate the antitumor activity of CDK4/6 inhibitors using *in vitro* and *in vivo* preclinical models of MPM. Based on publicly available transcriptomic data of MPM, patients with *CDK4* or *CDK6* overexpression had shorter overall survival. Treatment with abemaciclib or palbociclib at 100 nM significantly decreased cell proliferation in all cell models. Both CDK4/6 inhibitors significantly induced G1 cell cycle arrest thereby increasing cell senescence and increased the expression of interferon signaling pathway and tumor antigen presentation process in culture models of MPM. *In vivo* preclinical studies showed that palbociclib significantly reduced tumor growth and prolonged overall survival in a platinum-naïve and platinum resistant MPM mouse model. Treatment of MPM with CDK4/6 inhibitors decreased cell proliferation, mainly by promoting cell cycle arrest at G1 and by induction of cell senescence. Our preclinical studies provide evidence for evaluating CDK4/6 inhibitors in the clinic for the treatment of MPM.

## Introduction

Malignant pleural mesothelioma (MPM) is an aggressive, locally invasive and currently not curable malignancy of the pleura, which is associated with occupational and para-occupational exposure to asbestos (1). Although asbestos use is banned in many countries, asbestos-insulated buildings are present throughout the world and some countries are still manufacturing and using large quantities of asbestos (2).

Treatment options are limited for patients with advanced MPM (3). Palliative chemotherapy consisting of platinum and pemetrexed is the established standard of care for MPM in patients with advanced disease with ECOG performance status of 0-2 (4). The addition of bevacizumab to chemotherapy modestly improved overall survival, but this treatment is not available in all countries (5). To date, no treatment has yet been shown to improve survival in the relapsed setting, resulting in a high unmet need for effective therapies in previously treated patients with MPM. There are currently no approved targeted therapies for mesothelioma. Single agent immunotherapy has demonstrated limited efficacy in the relapsed setting in a randomized phase III clinical trial (6). Dual immune checkpoint inhibition of PD1 and CTLA-4 has recently demonstrated superiority to platinum plus pemetrexed in the first-line setting in the CheckMate-743 study and is likely to change the treatment landscape in MPM (7).

Characterization of the genomic landscape of MPM has revealed a high frequency of recurring focal and arm-level deletions, reflecting a predominant loss of tumor suppressor genes in MPM (8). Cyclin dependent kinase inhibitor 2A (*CDKN2A)* deletions are found in 56-70% of MPM and are associated with shorter overall survival (9,10). The C*DKN2A/ARF* locus (9p21) encodes for two cell cycle regulatory proteins: p14ARF and p16INK4a, the latter being a negative regulator of cyclin-dependent kinase 4/6 (CDK4/6) (11,12). In a recent clinical trial of personalized therapy in advanced NSCLC, *CDKN2A* loss was associated with sensitivity to CDK4/6 inhibitors (11,13). Considering the high frequency of *CDKN2A* deletions in MPM and the fact that cell cycle deregulation is a hallmark of this disease, we postulated that CDK4/6 inhibitors might constitute a novel therapeutic approach in MPM. In the last decade, several selective CDK4/6 inhibitors, abemaciclib, ribociclib and palbociclib, have been approved for the treatment of metastatic breast cancer (14-16).

In the present work, we report that *CDK4* or *CDK6* overexpression in primary tumors conferred a more aggressive behavior in patients with MPM based on publicly available transcriptomic data. We assessed the efficacy of CDK4/6 inhibitors both *in vitro* in MPM cell models and *in vivo* in subcutaneous and orthotopic xenografts to investigate their potential in the treatment of MPM.

## Results

### Overexpression of CDK4 and CDK6 are associated with poor prognosis in patients with MPM

Patients with *CDK4* overexpression (i.e., above the median) had significantly shorter overall survival in both cohorts (12.6 and 13.3 months, respectively) compared with patients with lower expression (23.5 and 25.9 months respectively). *CDK4* overexpression remained statistically significant after adjusting by age, gender, tumor stage and histologic subtype in each dataset, as well as in the combined analysis of both series (HR=2.10 [95% CI 1.53–2.88]; p=4.2e-06; **Figure 1A**). Patients with *CDK6* overexpression (i.e., above the median) had significantly shorter overall survival (12.6 months) compared with patients with lower expression (20.3 months; p=0.00026) in the Bueno cohort and there was a trend toward shorter overall survival in the TCGA dataset (15 versus 23.6 months; p=0.060). Nevertheless, *CDK6* overexpression remained statistically significant in the combined analysis after adjusting by age, gender, tumor stage and histologic subtype in the Bueno cohort, and also in the combined cohort (HR=1.74 [95% CI 1.32–2.29]; p=5.4e-05; **Figure 1B**).

**Figure 1.**
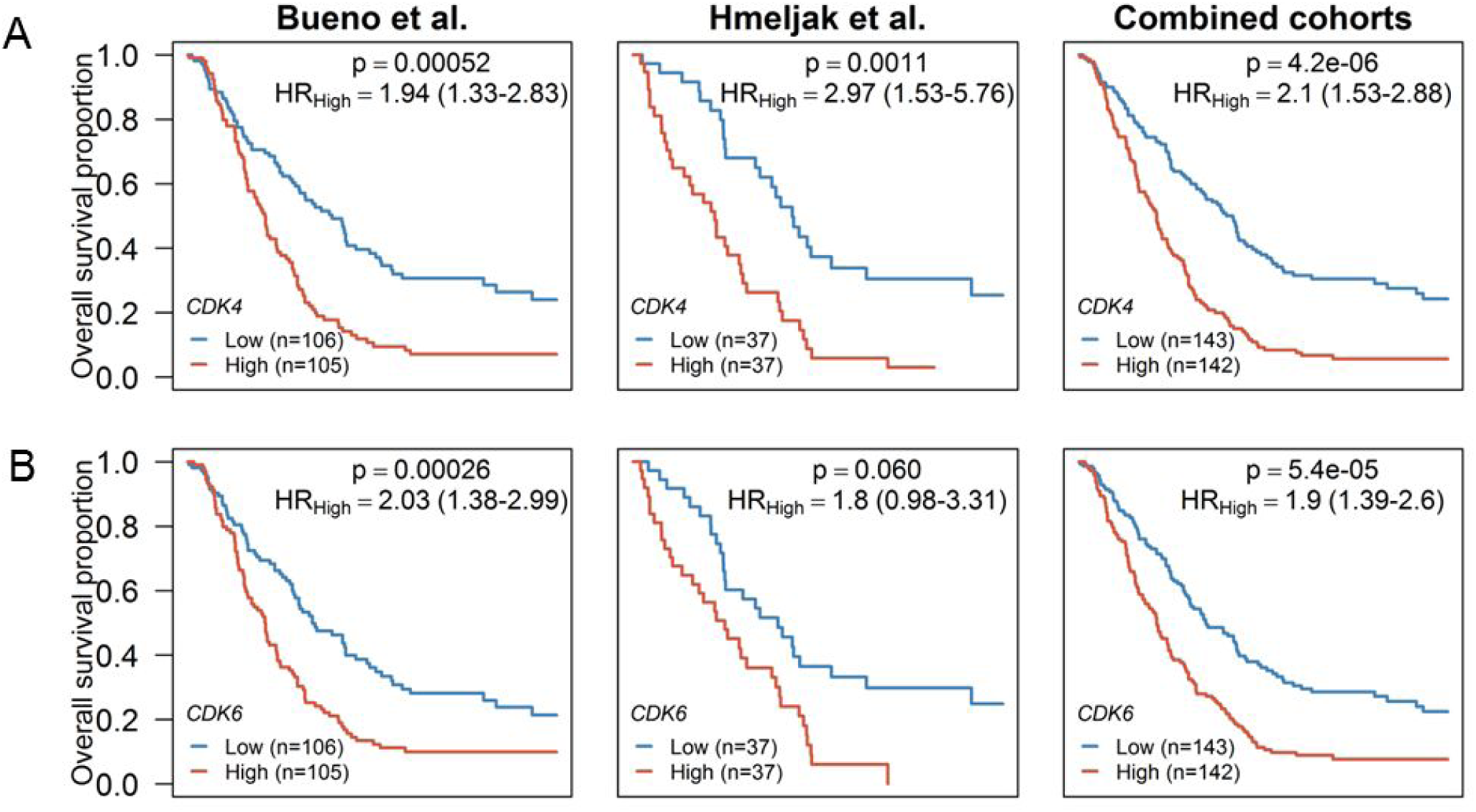
Kaplan-Meier plots of overall survival (OS) in MPM patients according to *(A) CDK4* and *(B*) *CDK6* gene expression levels based on data obtained from Bueno et al. (left column) and Hmeljak et al. (middle column) cohorts or the combination of both (right column). High levels of *CDK4* or *CDK6* (red line) were significantly associated with poor OS in patients with MPM. In each cohort, the high and low expression levels were defined based upon the median. P-values and hazard ratios (HR) were calculated by likelihood ratio test and multivariate Cox regression analysis respectively.

As previously reported (9,10), low expression of *CDKN2A* was associated with shorter overall survival in both cohorts and was independently associated with worse prognosis in the combined cohort including Bueno and TCGA (HR=0.49 [95% CI 0.36–0.66]; p=3.4e-06; **Supplementary Figure S1A**). In the TCGA cohort, only two tumors harbored an *RB1* homologous deletion, while *CDKN2A/p16* deletion was a common event present in 34 out of 74 cases (46%).

*CDKN2A* copy number was assessed in an independent cohort of 79 MPM acquired at radical surgery involving extended pleurectomy decortication. Patient clinicopathological characteristics are outlined in **Table 1**. Homozygous loss of 9p21.3 encompassing *CDKN2A* was observed in 40 samples (50.6%), while copy number loss/LOH was observed in 18 (22.7%). *CDKN2A* homozygous loss was associated with shorter median overall survival (10.98 months) compared to euploid *CDKN2A* (45.8 months; HR=0.37 [95% CI 0.22 - 0.62]; p=0.0002; **Supplementary Figure S1B**). *CDKN2A* copy number loss/LOH was associated with shorter median overall survival (8.52 months) compared to wild-type CDKN2A (45.8 months; HR=0.18 [95% CI 0.08-0.40]; p=0.0001). There were no statistically significant differences in overall survival among patients harboring *CDKN2A* homozygous deletion compared to those with *CDKN2A* copy number loss/LOH (HR=0.89 [95% CI 0.49-1.59]; p=0.158).

**Table 1.**
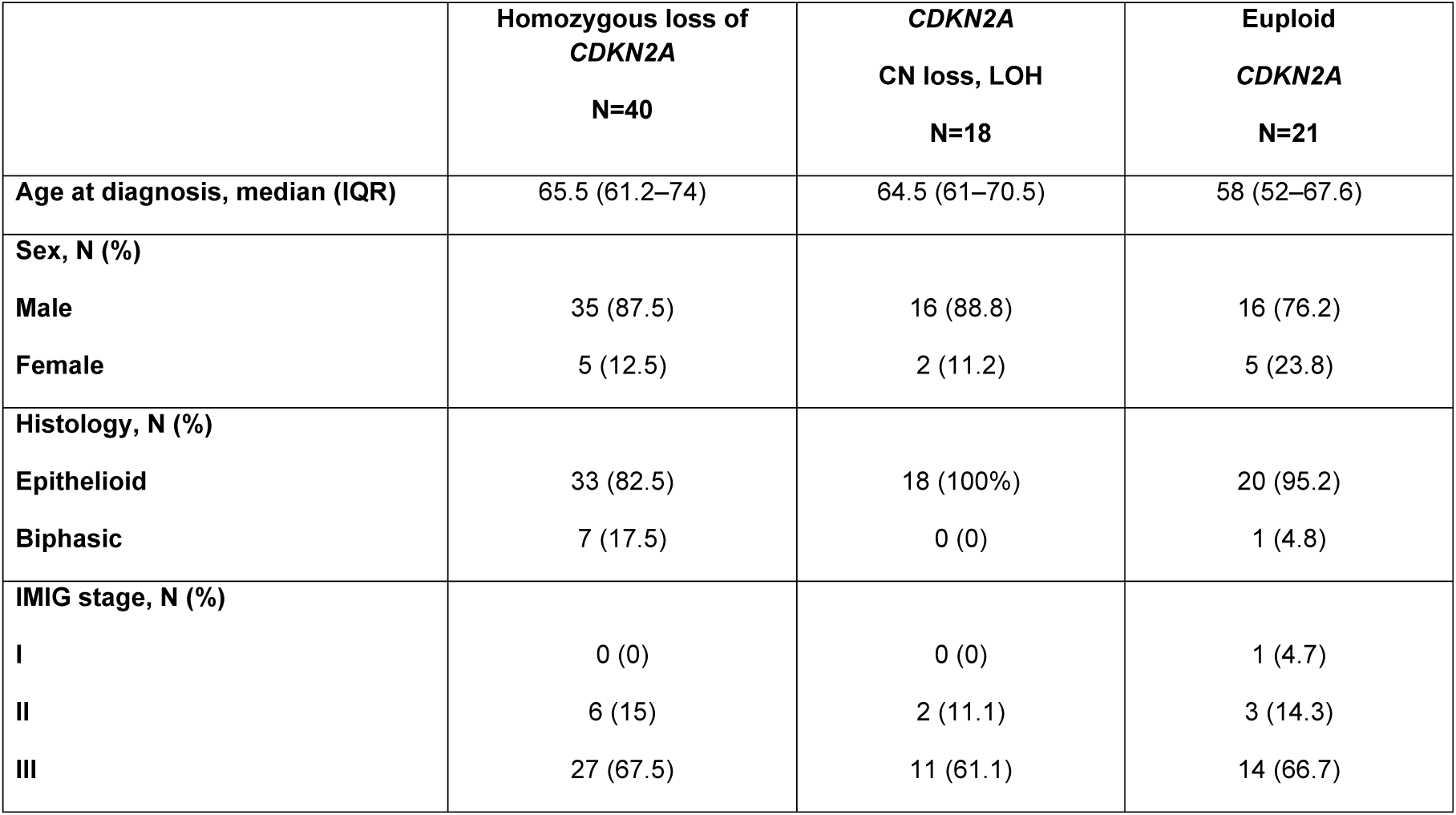
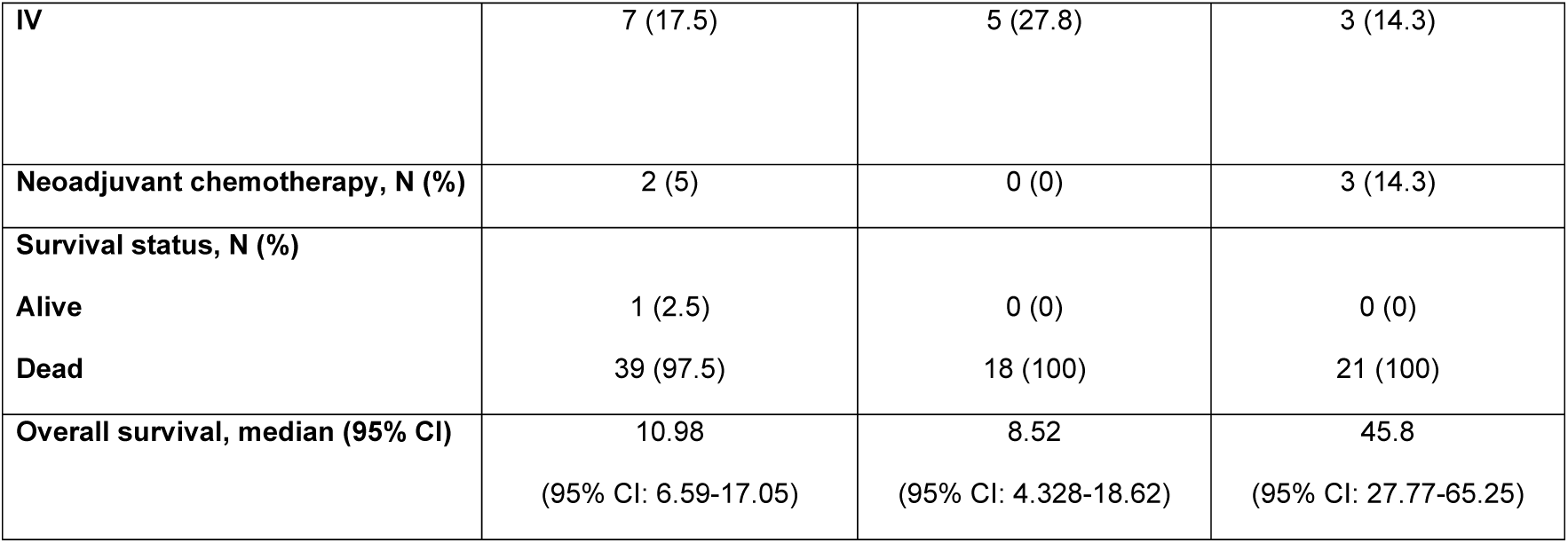
Clinicopathological characteristics of patients. (CI: confidence interval; CN: copy number; IMIG: International Mesothelioma Interest Group; IQR: interquatile range; LOH: loss of heterozygosity).

|  | **Homozygous loss of**<br>***CDKN2A***<br>**N=40** | ***CDKN2A***<br>**CN loss, LOH**<br>**N=18** | **Euploid**<br>***CDKN2A***<br>**N=21** |
| --- | --- | --- | --- |
| **Age at diagnosis, median (IQR)** | 65.5 (61.2–74) | 64.5 (61–70.5) | 58 (52–67.6) |
| **Sex, N (%)** |  |  |  |
| Male | 35 (87.5) | 16 (88.8) | 16 (76.2) |
| Female | 5 (12.5) | 2 (11.2) | 5 (23.8) |
| **Histology, N (%)** |  |  |  |
| Epithelioid | 33 (82.5) | 18 (100%) | 20 (95.2) |
| Biphasic | 7 (17.5) | 0 (0) | 1 (4.8) |
| **IMIG stage, N (%)** |  |  |  |
| I | 0 (0) | 0 (0) | 1 (4.7) |
| II | 6 (15) | 2 (11.1) | 3 (14.3) |
| III | 27 (67.5) | 11 (61.1) | 14 (66.7) |
| **IV** | 7 (17.5) | 5 (27.8) | 3 (14.3) |
| **Neoadjuvant chemotherapy, N (%)** | 2 (5) | 0 (0) | 3 (14.3) |
| **Survival status, N (%)** |  |  |  |
| **Alive** | 1 (2.5) | 0 (0) | 0 (0) |
| **Dead** | 39 (97.5) | 18 (100) | 21 (100) |
| **Overall survival, median (95% CI)** | 10.98<br>(95% CI: 6.59-17.05) | 8.52<br>(95% CI: 4.328-18.62) | 45.8<br>(95% CI: 27.77-65.25) |

### Genomic characterization of patient-derived MPM models and baseline expression of genes involved in cell cycle in MPM cell lines

Clinicopathological characteristics and main genomic alterations are shown in **Table 2**. In all three patient-derived cell lines, *CDKN2A/p16* was deleted and *NF2* was wild type, while *BAP1* was mutated in ICO_MPM1 (p.K651Yfs*1) and ICO_MPM2 (p.R60X). Additional information about their mutational profile is provided in **Supplementary Table S1**.

**Table 2.**
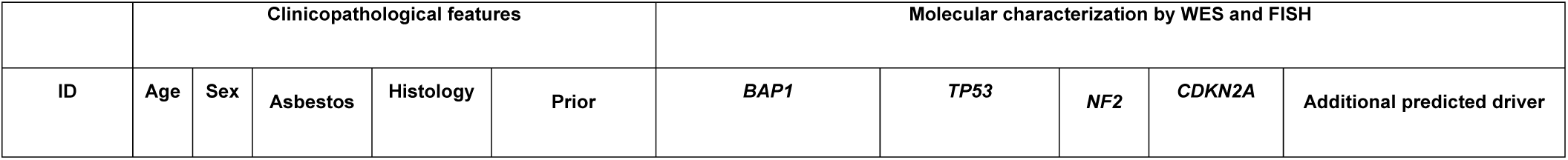
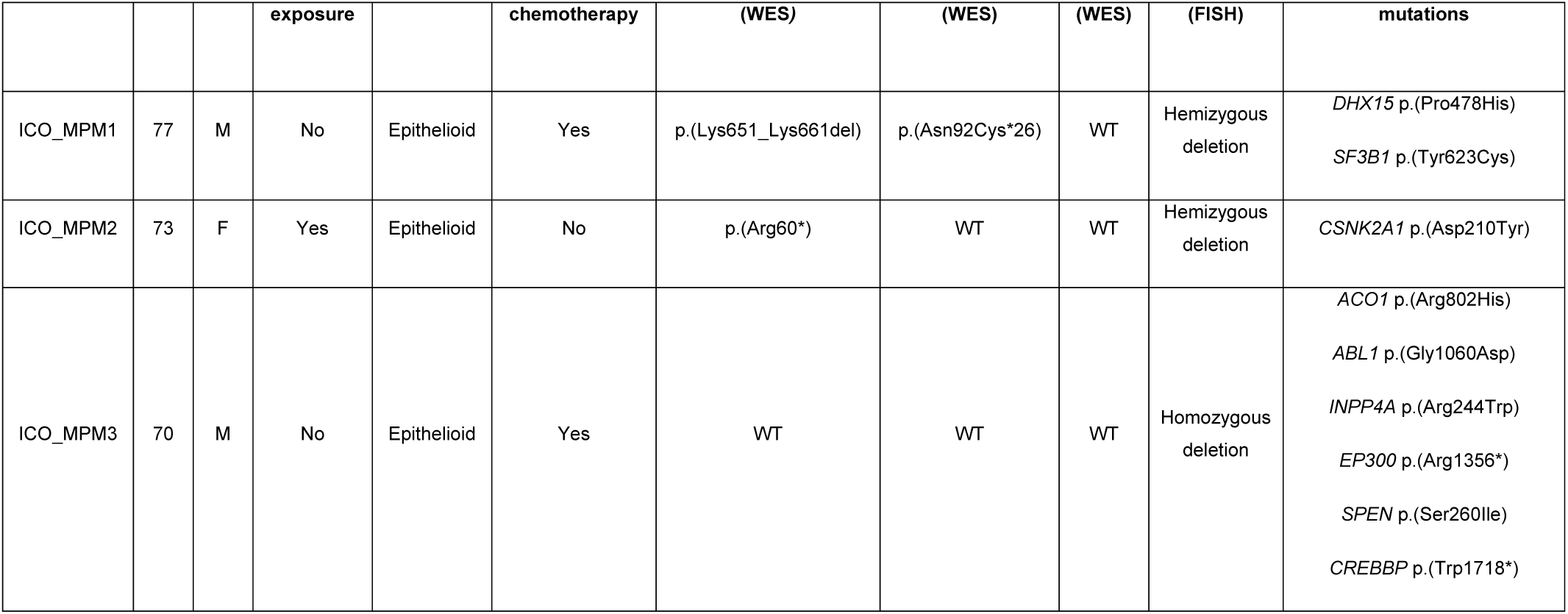
Clinicopathological characteristics and main genomic and protein alterations found in primary cell lines were derived from patients with pleural malignant mesothelioma. Additional predicted driver mutations have been identified using Cancer Genome Interpreter. (WES: whole exome sequencing; FISH: fluorescence in situ hybridization).

We examined by Western blot the expression levels of CDK4, CDK6, cyclin D1, CDKN2A/p16 and RB proteins in five commercial and in the three patient-derived MPM cell lines (**Figure 2A**). p16 expression was not detected in any cell line, while CDK4 was highly expressed in MSTO-211H, H28, H2052 and H2452 and in all primary cell lines. CDK4 expression was lower in H226, but levels were still perceptible. High CDK6 expression was detected in MSTO-211H, H28, H226 and ICO_MPM2 whereas its expression in H2052, H2452, ICO_MPM1 and ICO_MPM3 was lower. Five out of eight cell lines showed high expression of cyclin D1 and the other three expressed lower, but detectable levels. None of the cell lines showed loss of RB expression. Additional information about their mutational profile is provided in **Supplementary Table S2**.

**Figure 2.**
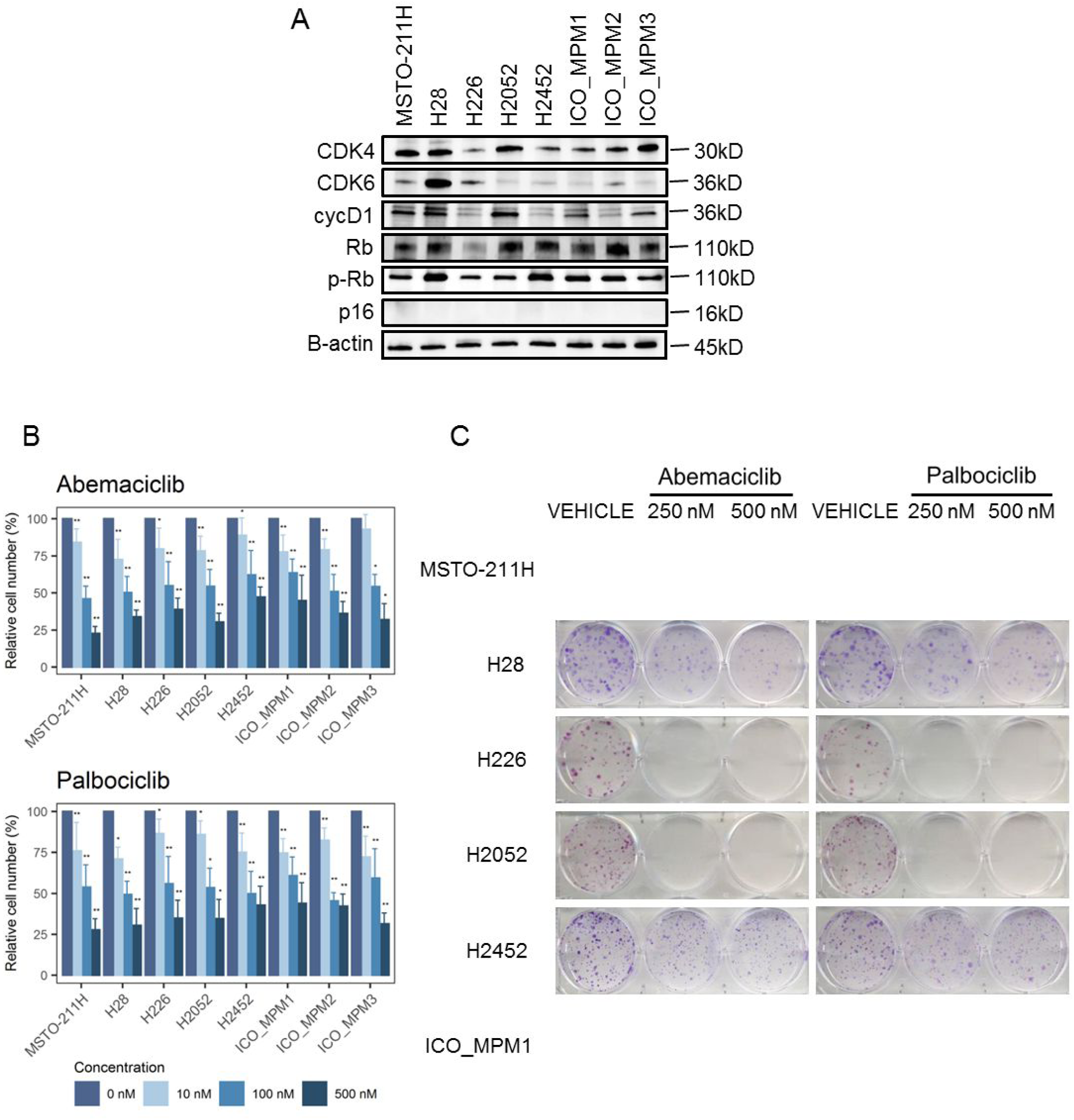
Quantification of the expression levels of key cell cycle regulators and response to treatment with CDK4/6 inhibitors in a panel of commercial MPM cell lines (MSTO-211H, H28, H226, H2052 H2452) and primary patient-derived cultures (ICO_MPM1, ICO_MPM2, ICO_MPM3). *(A)* Base line protein expression levels by Western blot of CDK4, CDK6, cyclin D1, Rb, phosphor-RB and p16. *(B)* Number of viable cells was determined *in vitro* by cell counting in the panel of cells after three days of treatment with increasing concentrations (0, 10, 100, 500 nM) of abemaciclib or palbociclib. Bar plots represent the means ± SD of three measurements. Adjusted p-values were calculated with Wilcoxon signed rank tests. In the graph, the p values are reported with respected to 0 nM (*p < 0.05; **p < 0.01). (*C*) Colony formation assay displaying treatment response to abemaciclib and palbociclib. A representative image from 3 biological independent replicates is displayed.

### Antiproliferative effect of CDK4/6 inhibitors on human MPM cell lines

All MPM cell lines treated with increasing concentrations of abemaciclib or palbociclib for 72 hours, showed a decrease in cell number (**Figure 2B and Supplementary Figure S2)**. Treatment with abemaciclib and palbociclib at 100 or 500 nM significantly reduced cell number in comparison to control in all cell lines tested (p<0.05). At lower doses (10 nM), the decrease in cell number was statistically significant in six out of eight cell lines after treatment with abemaciclib (p<0.01), while it was significantly reduced in all cell lines after palbociclib treatment (p<0.05).

The reduction in cell number after exposure to CDK4/6 inhibitors at 100 nM was nearly 50% (mean 54.5% ± 5.5 with abemaciclib and mean 53.4% ± 4.9 with palbociclib). At 500 nM, a reduction of 64.3% and 64.1% was observed with abemaciclib and palbociclib respectively. MSTO-211H was the most sensitive cell line to both CDK4/6 inhibitors at 100 nM and 500 nM doses. All primary cell lines were sensitive to CDK4/6 inhibitors regardless of whether they had been derived from a patient who was chemotherapy-naïve or who had received prior chemotherapy. ICO_MPM2, which was derived from a chemotherapy-naïve patient, was the most sensitive primary cell line to palbociclib with a cell number reduction of 45.3% ± 5.2 at 100 nM (**Figure 2B**). These antiproliferative effects were confirmed by cell colony formation assay and crystal violet staining (data not shown). The ability to form colonies was completely blocked when MSTO-211H, H226, H2052 and ICO_MPM1 were treated with abemaciclib and palbociclib at 250 and 500 nM **(Figure 2C)**.

### Effect of CDK4/6 inhibitors on cell cycle in human MPM cell lines

Three cell lines selected for expressing high levels of CDK4 and CDK6 proteins (MSTO-211H, H28 and ICO_MPM3) were evaluated for alterations in cell cycle progression after 24-hour treatment with 250 or 500 nM abemaciclib or palbociclib. Compared with control, cells treated with abemaciclib or palbociclib were arrested at G1 phase (**Figure 3A**).

**Figure 3.**
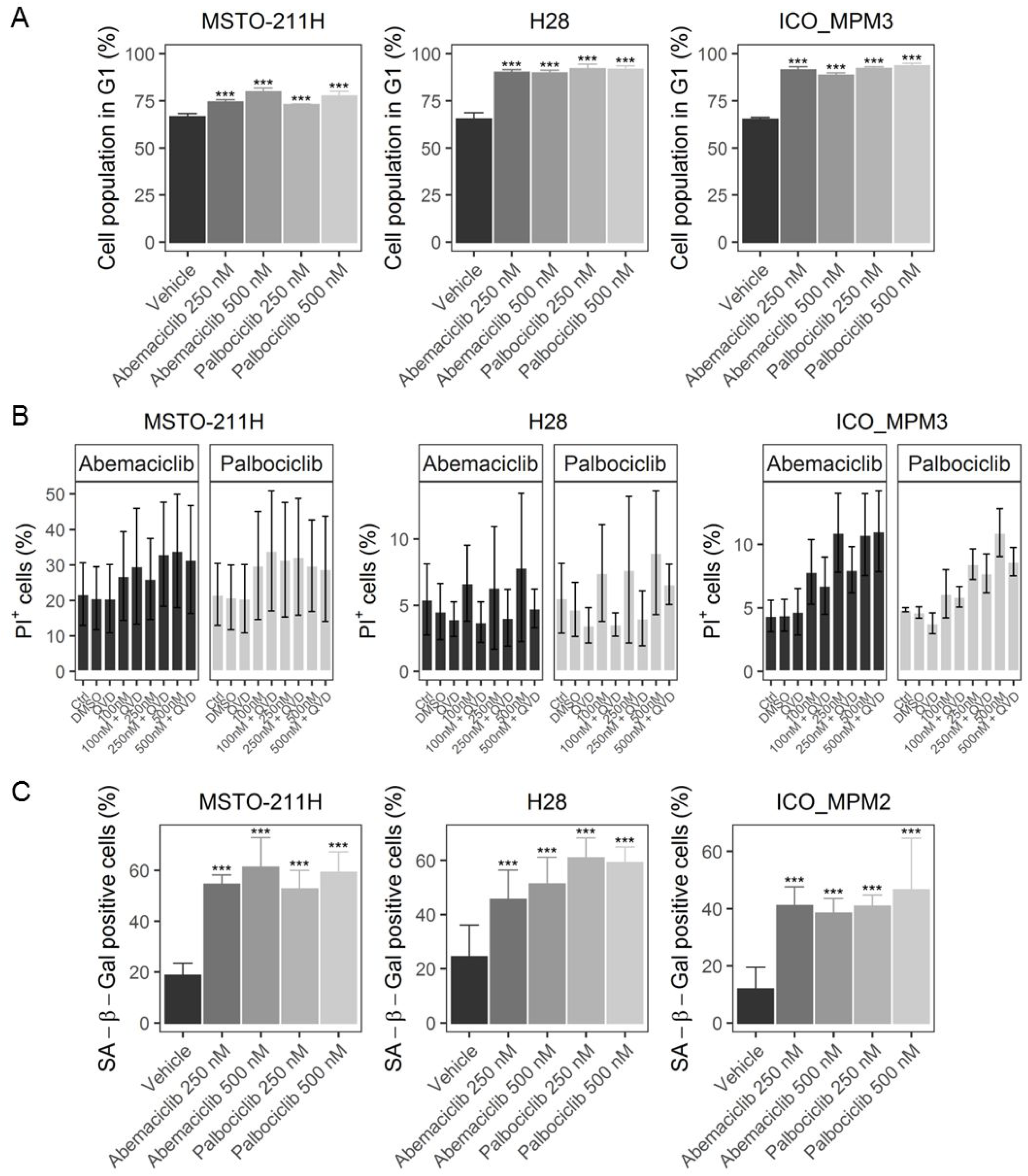
Effects of cell line treatments with CDK4/6 inhibitors abemaciclib or palbociclib at 0, 100, 250 or 500 nM doses to induce *(A)* cell-cycle arrest (*B*), cell death and (*C*) senescence. (*A*) MSTO-211H, H28 and ICO_MPM3 cells were untreated and treated with both inhibitors during 24 hours and DNA content analyzed by flow cytometry. Cell cycle arrest at G1 phase was induced by both CDK4/6 inhibitors in the cell lines. (*B*) Likewise the same three cells were exposed to drugs for 72 hours and cell death stained with propidium iodide (PI^+^ cells) analyzed by flow cytometry. Cell death was slightly affected (up to 1-5%) by both drugs. To quantify the percentage of non-apoptotic induced cell death, cells were treated with QVD (an apoptosis inhibitor). Cell cycle phase distribution and cell death analysis were done using FlowJo software. (*C*) A significant increase in the number of SA-β-Gal positive cells was detected in MSTO-211H, H28 and ICO_MPM2 treated. Data are expressed as a percentage of senescent cells obtained from the mean value ± SD of three replicates. Adjusted p-values less than 0.05 were considered significant.

At 500 nM, the fraction of cells at G0/G1 phase increased to 13% and 11% in MSTO-211H, to 24% and 26% in H28 and to 23% and 28% in ICO_MPM3 after abemaciclib or palbociclib treatment, respectively (p<0.001). In addition, a significant decrease in cell percentage in the G2/M phase (p<0.001) and in S phase (p<0.001) was observed in the three cell lines after treatments at 250 nM and 500 nM compared to non-treated cells (**Supplementary Figure S3**). The percentage of subG1 fraction significantly increased from 1.38% in MSTO-211H control cells to 2.32% ± 0.6 after treatment (p<0.001, **Supplementary Figure S3**). In H28, the percentage of subG1 fraction significantly decreased from 3.36% to 1.63% ± 0.2 after treatment (p<0.001, **Supplementary Figure S3**). In ICO_MPM3, the percentage of cells in subG1 remained unchanged from 0.90 in control cells to 0.94% ± 0.1 under all treatments except with 250 nM palbociclib, where there was a small, but statistically significant increase (0.9% vs 1.07%; p<0.05, **Supplementary Figure S3**). These small biological differences in cells with subG1 DNA content suggest that there is no increment in classical apoptosis after abemaciclib or palbociclib at different exposure concentrations.

### Effect of CDK4/6 inhibitors on cell death and senescence in human MPM cell lines

As a next step, we investigated whether treatment with CDK4/6 inhibitors could induce cell death in MPM cells. MSTO-211H, H28 and ICO_MPM3 cells were treated with different concentrations of abemaciclib and palbociclib, as single agents or in combination with the apoptosis inhibitor QVD for 72 hours and cell death was quantified by FACS (**Figure 3B**). Neither inhibitor was able to significantly increase the levels of apoptosis in MSTO-211H, H28 and ICO_MPM3 cells at any of the doses tested. At the highest dose (500 nM), the percentage of apoptotic cells reached 12% with abemaciclib and 8% with palbociclib in MSTO-211H cells, 2% after abemaciclib and 3% after palbociclib in H28 cells, and around 6% after either treatment in ICO_MPM3 cells.

To investigate whether CDK4/6 inhibitors promote senescence, both treated and control MSTO-211H, H28 and ICO_MPM2 cells were stained using β-galactosidase. A significant increase in the percentage of senescent cells was detected in all cell lines treated with different concentrations of abemaciclib or palbociclib (p<0.001, **Figure 3C**). Specifically, the proportion of senescent SA-β-gal positive MSTO-211H cells increased from 18% to 54% with 250 nM abemaciclib and to 61% with 500 nM abemaciclib and to 52% with 250 nM palbociclib and to 59% with 500 nM palbociclib. Likewise, an increase of senescent cells was also observed in H28 cells treated with 250 nM or 500 nM of either inhibitor. In ICO_MPM2 cells, the percentage of SA-β-gal positive cells increased from 12% in control cells to 41% after abemaciclib and to 46% after palbociclib treatments at 250 and 500 nM, respectively.

### Gene-expression profiling in cell lines and xenografts treated with CDK4/6 inhibitors

To determine the functional consequences of CDK4/6 inhibitor treatment, we performed transcriptomic analysis of MSTO-211H cells treated with abemaciclib or palbociclib at 250 nM for 72 hours. In addition, tumor xenografts treated with palbociclib were also evaluated. After treatment with either CDK4/6 inhibitor, a significant downregulation in the expression levels was observed in the MSTO-211H cell line for genes related with cell cycle, such as regulation of transcription genes involved in G1-S transition of mitotic cell cycle, nucleus organization and mitotic spindle assembly and organization (**Supplementary Figure S4 and Supplementary Table S3**). On the other hand, there was a significant upregulation of genes related to interferon signaling pathways, lymphocyte migration and chemotaxis, complement activation and antigen presentation pathways, such as MHC protein complex. Furthermore, the transcriptomic analysis of palbociclib-treated tumors xenografts mice showed similar results (data not shown).

### Palbociclib reduced tumor growth in *in vivo* preclinical tumor models improving overall survival in mice with orthotopically implanted MPM tumors

The effect of palbociclib *in vivo* was examined by implanting subcutaneously MSTO-211H cells into the right flanks of athymic mice. After 26 days of treatment, the mean volume of tumors implanted subcutaneously in vehicle-treated mice was 1816 ± 795.2 mm^3^; in cisplatin plus pemetrexed-treated mice was 1647.1 ± 733.8 mm^3^ whereas for palbociclib-treated mice it was 524.2 ± 236.6 mm^3^ (**Figure 4A and Supplementary Figure S5A**). Differences among palbociclib and the two other cohorts were already statistically significant at day 16 (p=0.043, **Supplementary Figure S5B**). At mice sacrifice, 26 days post-treatment, a significant decrease in the tumor weight was observed for palbociblib-treated mice respect to vehicle and combined chemotherapy-treated mice (0.35 vs 1.1 and 1.13 gr; p=0.01 and p=0.007, respectively, **Figure 4B and 4C**). No differences were observed at histologic level **(Supplementary Figure S6)**. The body weight of the mice was monitored to evaluate the potential side effects of treatments (**Supplementary Figure S5C**). In those mice treated with palbociclib, no body weight loss was observed during the experiment, suggesting that palbociclib did not exert significant systemic toxicity at the doses used in this study.

**Figure 4.**
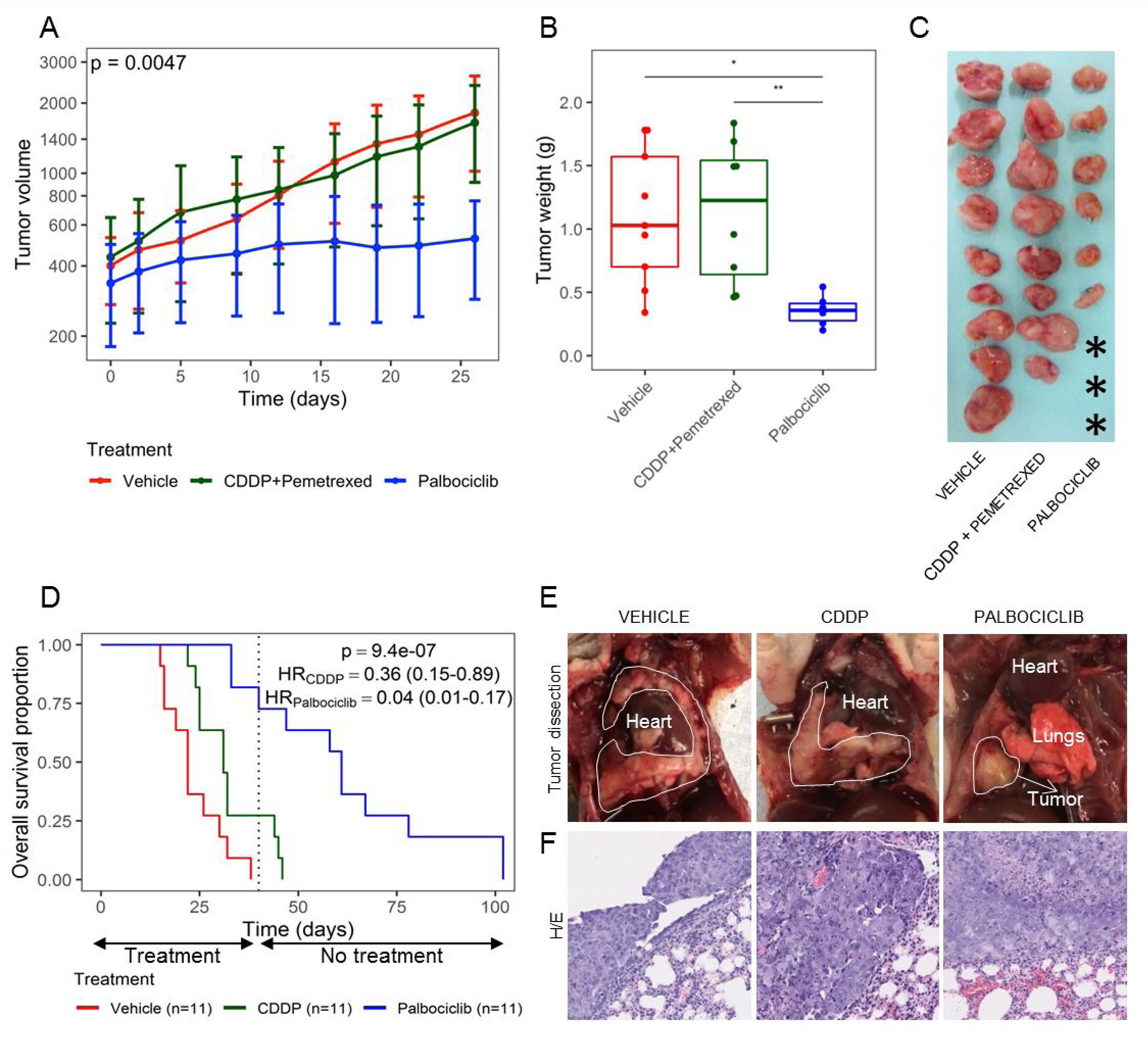
*In vivo* treatment with palbociclib in both subcutaneous and advanced orthotopic MPM models. A xenografted subcutaneous tumor model was establishedby inoculation of MSTO-211H cells into the flanks of athymic nude mice. Tumors’+ volume (*A*) was monitored by caliper measure every four days, and at the end of the experiment, mice were sacrificed and *(B* and *C)* the tumors were removed, weighted and photographed. An Advanced orthotopic model was generated by implantation in the lung of mice (n=35) of small solid fragments (2-3 mm^3^) of previously generated MSTO-211H subcutaneous cisplatin plus pemetrexed resistant tumor xenograft. (*D*) Kaplan-Meier curves showing survival of MSTO-211H orthotopic tumor-bearing mice. (*E*) Representative MSTO-211H images of orthotopic tumors dissected from each group of treatment and *(F)* histological characterization on H&E sections (Scale bars = 100 µm). Orthotopic model accurately reproduce human MPM disease characteristics as tumor grown from the site of implantation to all the pleural space. Tumor mass area is delimited by white line. Asterisks indicated absence of apparent macroscopic tumor at sacrifice, while residual cells were identified by H&E analysis.

To preclinically investigate the efficacy of palbociclib as second-line treatment in MPM tumors refractory to conventional chemotherapy, we re-implanted one of the subcutaneous tumors derived from MSTO-211H cells previously treated with cisplatin plus pemetrexed. After 40 days of treatment, no vehicle-treated mice were alive (0/11; 0%); only three platinum-treated mice were alive (3/11; 27.3%), while seven palbociclib-treated mice were still alive (7/11; 63.6%) (**Figure 4D**). Animals (93.9%) were sacrificed due to dyspnea or excessive weight loss (31/33). Overall survival analysis showed a significant reduction in the risk of death for palbociclib-treated mice compared with vehicle (HR=0.04 [95% CI 0.01-0.17]) or with cisplatin (HR=0.11 [95% CI 0.03-0.41]).

Those mice that were alive after 40 days of treatment (n=10) were maintained without treatment and followed up until endpoint. After six days of stopping treatments, all the remaining platinum-treated mice (n=3) were dead, whereas two out of seven palbociclib-treated mice were still alive after two months without receiving any treatment. Palbociclib treatment did not exert any substantial change in body weight between the first and the last day of treatment **(Supplementary Figure S5D)**. Representative pictures of MSTO-211H orthotopic tumors from each group of treatment are shown in **Figure 4E and Supplementary Figure S7**. Histopathological analysis of the MPM tumor xenografts grown orthotopically in mice accurately reproduced the natural history of mesothelioma **(Figure 4F and Supplementary Figure S8)**.

## Discussion

MPM is a rapidly fatal neoplastic disease in which therapeutic options are limited and there are no effective second-line treatment available. We investigated the role of CDK4/6 inhibition in MPM because cell cycle deregulation is a relevant hallmark in this disease. In this regard, *CDKN2A/p16* deletion is a common genomic event associated to worse clinical outcome in MPM (9,10). Based on publicly available gene expression data of MPM, we found that overexpression of *CDK4* or *CDK6* is associated with shorter overall survival. The unfavorable prognostic role of *CDKN2A/p16* deletion was confirmed in an independent cohort of MPM. Together these findings suggest that cell cycle deregulation may confer an aggressive biological behavior in mesothelioma and let us to hypothesize that treatment with CDK4/6 inhibitors might be an effective treatment in MPM.

The efficacy of palbociclib has been previously studied in *in vitro* models of MPM (17). However, the antitumor activity of CDK4/6 inhibitors has not yet been evaluated using primary patient-derived cell models of MPM neither *in vivo* models of MPM. In our work, we assessed the efficacy of two CDK4/6 inhibitors, abemaciclib and palbociclib, in a subset of five commercial and three primary patient-derived cell culture models obtained from pleural effusions of patients with MPM (one chemotherapy-naïve and two after progression to standard first-line chemotherapy). Furthermore, we performed not only *in vivo* basic subcutaneous drug response studies in xenografts derived from one chemotherapy-naïve MPM cell line, but also advanced studies by means of orthotopic implanted xenograft from a cell line-derived tumor previously treated with cisplatin plus pemetrexed. Remarkably all the cell lines were sensitive to palbociclib and to abemaciclib. Treatment with abemaciclib or palbociclib significantly reduced cell proliferation, as evaluated by cell number counting or by colony formation ability, in all the cell lines including in the primary ones derived from pleural liquid from patients resistant to chemotherapy. Interestingly, these *in vitro* experiments underscore two important points: i) no substantial differences were found in the antiproliferative effect of both inhibitors; and ii) the sensitivity to CDK4/6 inhibitors was not correlated with the endogenous expression levels of CDK4 or CDK6.

Then, we assessed the impact of abemaciclib and palbociclib treatment on cell cycle progression, cellular senescence and apoptosis induction. These experiments were performed using three cell lines that were sensitive to both drugs, including a primary cell line derived from a patient resistant to chemotherapy. As expected, the treatment with abemaciclib and palbociclib caused cell cycle arrest at G1 phase but also promoted cellular senescence. However, neither abemaciclib nor palbociclib activated programmed cell death or apoptosis as indicated by negligible subG1 accumulation or propidium iodide incorporation. The observed increase on cellular senescence induced by both drugs could be linked to apoptosis resistance mechanisms (15,18). As proposed by other groups (17,19,20), our results reinforce the cytostatic mechanism of action of CDK4/6 inhibitors and underscore that these should be given sequentially after completing chemotherapy treatment (21). However, the low cell death observed *in vitro* does not eliminate the possibility that palbociclib, by inducing senescence in a few cells, may promote *in vivo* cytotoxicity mediated by Natural Killer cells. This phenomenon has been described in lung cancer, in which Natural Killers participate in tumor reduction upon treatment with MEK inhibitors and palbociclib (22,23).

In order to explore potential activation of compensatory pathways, we performed gene set enrichment analysis of the transcriptome of MSTO-211H cells treated with abemaciclib or palbociclib *in vitro* or subcutaneously implanted in mice. This experiment showed downregulation of genes involved in G1-S transition of mitotic cell cycle, nucleus organization and mitotic spindle assembly and organization. In concordance with studies conducted in other tumor types, genes encoding interferon signaling and antigen presentation pathways were upregulated after CDK4/6 pharmacological inhibition (20). In melanoma, CDK4/6 inhibition activates p53 by lowering PRMT5 which leads to altered MDM4 splicing and significantly reduced protein expression (24,25). Other studies performed in breast cancer cell lines and transgenic mice models have shown that abemaciclib treatment increased the expression of antigen processing and presentation and even suppressed the proliferation of regulatory T cells (16,20). Further studies evaluating the functional consequences of the treatment with CDK4/6 inhibitors on the tumor immune contexture are warranted in mesothelioma.

As both CDK4/6 inhibitors showed a similar pattern of response and effectiveness *in vitro* and considering that palbociclib has a more convenient posology compared with abemaciclib, we decided to perform *in vivo* experiments with palbociclib. To the best of our knowledge, this is the first time that effectiveness of palbociclib was evaluated using preclinical *in vivo* subcutaneous and orthotopic MPM tumor models. Among the available cell line models, MSTO-211H cell line was selected for the *in vivo* experiments because i) it expresses CDK4 and CDK6; ii) it is the most sensitive cell line to palbociclib at 500 nM; and iii) it was tumorigenic in athymic mice. Our results showed that palbociclib reduced tumor size in a subcutaneous mouse model of chemo-naïve MSTO-211H cells compared with standard chemotherapy (cisplatin plus pemetrexed). An increased expression of pro-apoptotic proteins after long-term chemotherapy treatment could explain the chemoresistance in this model (26). We tried to replicate a situation representing treatment after progression to platinum-based chemotherapy by generating an orthotopic tumor mouse model and implanting in the pleura small solid fragments of MSTO-211H xenografted tumors previously treated with chemotherapy. In this advanced model of MPM, palbociclib significantly increased the overall survival of mice compared with cisplatin-based chemotherapy or vehicle; the benefits from treatment persisted even after stopping the treatment. Our results reinforce the potential use of palbociclib as a second-line treatment for patients with MPM that is resistant or has relapsed after standard chemotherapy doublet treatment. Interestingly, the extended response of palbociclib treatment, even after stopping treatment, could result from an effect modulating on the immune system response; however, this still needs to be elucidated.

Some limitations of our study are the absence of a wide range of available commercial MPM cell lines and the need for preclinical *in vivo* models representing the heterogeneity of the disease. However, in our study we have combined commercial, primary patient-derived lines as well as orthotopic models where mesothelioma grows in its corresponding microenvironment and can recapitulate the disease behavior.

A phase II clinical study of abemaciclib in patients harboring p16ink4a deficient, relapsed MPM has recently completed accrual (NCT03654833). CDK4/6 inhibition in this cohort has been associated radiological responses however the underlying molecular correlates of response are under investigation. Accordingly, whole exome sequencing of the trial cohort is planned to uncover genomic determinants of response.

In conclusion, our data support that treatment with CDK4/6 inhibitors, abemaciclib or palbociclib, can reduce cell proliferation and induce cellular senescence in MPM cell lines and palbociclib can increase overall survival of mice with orthotopically implanted MPM cells. A remarkable and sustained response to palbociclib was observed in xenografts of MPM tumor resistant to cisplatin and pemetrexed which was then implanted orthotopically in the pleural space of mice. Transcriptomic analysis of cell lines and xenografted tumors treated with CDK4/6 inhibitors showed an increased expression of interferon signaling pathway and antigen presenting processes, suggesting that CDK4/6 inhibitors may favor potential response to immunotherapy. Our results warrant further evaluation of CDK4/6 inhibitors as a second-line treatment in patients with advanced MPM that has failed standard platinum-based chemotherapy.

## Materials and methods

### Cell culture and cell lines

Five human MPM cell lines, including H28, H2452, H2052, MSTO-211H and H226 were purchased from the American Type Culture Collection (ATCC, Manassas, Virginia). Three additional primary cell lines (ICO_MPM1, ICO_MPM2 and ICO_MPM3) were derived from pleural effusions obtained from three patients with MPM. ICO_MPM1 and ICO_MPM3, were derived from two patients who progressed to standard chemotherapy with platinum and pemetrexed, while ICO_MPM2 was derived from a chemotherapy-naïve patient. Primary cells were isolated and cultured as previously described (27). All cell lines were incubated and maintained at 37°C in a humidified chamber containing 5% CO_2_.

### Patient and tissue samples

Patients with confirmed histological diagnosis of malignant pleural mesothelioma were scheduled for routine surgery involving extended pleurectomy decortication at the Glenfield Hospital (University of Leicester). Patients were approached 24 hours prior to their operation and provided with patient information regarding the research. All patients signed informed consent prior to surgery. Seventy-nine patient MPM samples were obtained at the time of surgery. Following surgery, all patients were longitudinally tracked until disease progression with CT monitoring, and monitored for survival.

### Oncoscan Analysis

DNA was extracted with the GeneRead DNA FFPE kit (Qiagen, Manchester, UK). Eighty nanograms of genomic DNA were analyzed using the OncoScan FFPE Assay Kit (Affymetrix, Wooburn Green High Wycombe, UK). The BioDiscovery Nexus Express 10.0 for OncoScan software was used to determine copy number alterations and loss of heterozygosity (LOH).

### Antibodies and drugs

Antibodies against total Rb (#9313), p-Rb (#8180), CDK4 (#12790), CDK6 (#13331), cyclin D1 (#2922), p16 (#80772) and β-actin (#4970) were purchased from Cell Signaling Technology (Danvers, Massachusetts) and were used following manufacturer instructions for western blot.

Abemaciclib (LY2835219) was purchased from Selleckchem (Houston, Texas). Palbociclib (PD0332991) was provided by Pfizer, Inc (San Diego, California). Cisplatin and pemetrexed were obtained at the Catalan Institute of Oncology pharmacy.

### *In vitro* and *in vivo* drug experiments

For *in vitro* experiments, cell lines were plated into 6-well plates and treated with abemaciclib or palbociclib with 0 (control), 10, 100, 250 or 500 nM for 1, 3 or 15 days. Doses below micromolar range, would be clinically well tolerated. For *in vivo* assays, mice were randomly treated with i) vehicle, 200 μl of 0.05 N sodium lactate pH 4.0 five days a week; ii) cisplatin alone, 3.5 mg/kg once a week or combined with pemetrexed, 100 mg/kg twice a week; or iii) palbociclib, 150 mg/kg five days over seven days. Mice were treated during twenty-six days for subcutaneous models or forty days for orthotopic models.

### Western blot analysis

Total cell lysates and western blotting were performed as previously described (28).

### Cell viability, cell cycle and apoptosis analysis

Cell viability was evaluated by cell counting and colony formation assays as described elsewhere (29). Cell cycle and apoptosis were analyzed as described in (30). A minimum of 1×10^4^ cells were analyzed per determination. All experiments were repeated at least three times with similar results. P-values were adjusted usingFDR.

### Measurement of cellular senescence

The evaluation of senescence-associated β-Galactosidase (SA-β-Gal) expression was performed as previously described (17). Experiments were repeated at least three times with similar results.

### *In vivo* MPM subcutaneous preclinical drug assays in nude mice

To investigate the efficacy of palbociclib in the treatment of MPM, we used the MSTO-211H cell line, derived from a patient who had not received prior chemotherapy and able to grow in athymic mice. For subcutaneous xenograft development, 4 × 10^6^ MSTO-211H cells growing exponentially were suspended in 300 µl PBS and subcutaneously inoculated into the right flanks of 30 four-week-old male athymic nude mice (Envigo, Indiana, Indianapolis). Once the tumors reached a homogeneous average volume size of 300–400 mm^3^, mice (n=28) were randomly assigned into four groups (n=7 per group) and treated as described above. To evaluate efficacy, tumor volumes (V=π/6 x L x W^2^) were measured twice per week with calipers and the weight of each animal was measured every day. After 26 days of treatment, mice were euthanized by cervical dislocation and the tumors were excised, weighted and processed for histologic and RNA studies following standard protocols. The mean volume + SD were calculated using R software v.3.5.0 (31). Daily differences among treatments were analyzed using Kruskal-Wallis tests, with FDR adjustment.

### *In vivo* MPM orthotopic preclinical drug assays in tumors nude mice

To investigate the efficacy of palbociclib in tumors after progression to standard first-line chemotherapy, two subcutaneous tumors treated with cisplatin plus pemetrexed from the previous experiments were aseptically isolated and implanted in 35 four-week-old male athymic nude mice following our previously reported procedures (32). Thirty-three mice were randomized into three groups (n=11 per group) and treated as previously mentioned for forty days. Beyond 40 days, all live mice remained untreated until human endpoint. Orthotopic tumors were collected from euthanized mice presenting breathing problems and processed for histological studies. Survival curves for each cohort of mice were calculated using the Kaplan-Meier method and the differences between groups were compared using Cox proportional hazards model.

### *In silico* analysis of publicly available RNA-sequencing data

Public data from RNA-seq cohorts published by Bueno et al. (33) and The Cancer Genome Atlas (TCGA-MESO) (8) were used to assess differences in survival. Gene expression (log_2_(TPM)) was stratified using the median, and Cox proportional-hazards models adjusted for sex, stage, age and histology were fitted to assess differences in survival using R software (31).

### Whole Exome Sequencing (WES) and RNA sequencing (RNA-seq) analysis of patient-derived cell lines

Paired-end RNA sequencing was performed on an Illumina HiSeq 2500, with 100 bp long reads. Genomic DNA and total RNA were submitted to the Centro National de Análisis Genómico (CNAG, Barcelona, Spain), for WES and RNA-Seq library preparation and sequencing. All statistical analyses were done using R software v.3.5.0 (31).

### Statistics

Cell proliferation assay was assessed using Wilcoxon signed rank tests comparing each treatment with vehicle condition, and adjusted using FDR correction. Differences among treatment and vehicle conditions in the cell death experiment were evaluated using Mann-Whitney U test for each comparison and Kruskal-Wallis test if 3 conditions were simultaneously tested and adjusted afterward using FDR. Cell cycle and senescence experiments were analyzed using proportion tests taking into account all the cells counted in the abovementioned experiments. P-values were adjusted using FDR. For *in vivo* experiments, the analysis of differences in body weight for orthotopic xenografts was computed using a Mann-Whitney U test comparing the first and last day of treatment in each treatment and adjusted with FDR. For subcutaneous xenografts, a longitudinal analysis testing for differences in treatment slopes was done using analysis of covariance. Regarding survival analysis, for public data (Bueno et al. (33) and The Cancer Genome Atlas (8), Cox proportional-hazards models adjusted for sex, stage, age, and histology were fitted to assess the differences between gene expression (categorized using the median log2TPM value for each gene). For *in vivo* experiments, a Cox proportional-hazards model was fitted to assess differences among groups. Survival curves were plotted using Kaplan-Meier curves. P-value smaller than 0.05 was considered statistically significant. All statistical analyses were done using R software v.3.5.0 (22).

### Ethics approval

The retrospective cohort was approved by a National Ethical Committee, under the references 4/LO/1527 (a translational research platform entitled *Predicting Drug and Radiation Sensitivity in Thoracic Cancers – also approved by University Hospitals of Leicester NHS Trust under the reference IRAS131283*) and 14/EM/1159 (retrospective cohort). Pleural effusions samples were obtained after patients signed the informed consent approved by the Hospital de Bellvitge Ethical Committee (PR152/14). All the animal experiments were performed in accordance with protocols approved by Animal Research Ethics Committee at IDIBELL. This study was performed in accordance with the principles outlined in the Declaration of Helsinki.

For additional information about methodology see supplementary material.

## Acknowledgements

We thank CERCA Program / Generalitat de Catalunya for their institutional support and grant 2017SGR448. E.Nadal received support from the SLT006/17/00127 grant, funded by the Department of Health of the Generalitat de Catalunya by the call “Acció instrumental d’+intensificació de professionals de la salut”.

## Author contribution statement

**EA:** Conceptualization, Methodology, Validation, Formal analysis, Investigation, Resources, Data Curation, Writing – Original Draft, Writing – Review & Editing, Visualization **AA:** Conceptualization, Software, Validation, Formal analysis, Resources, Data Curation, Writing – Review & Editing, Visualization **MMI**: Methodology, Investigation, Resources, Data curation **MHM:** Methodology, Validation, Investigation, Data Curation **DC:** Conceptualization, Software, Validation, Formal analysis, Data Curation **MG:** Conceptualization, Resources, Writing – Review & Editing **EP:** Resources, Writing – Review & Editing **MS:** Resources, Writing – Review & Editing **SB:** Software, Validation, Data Curation **AJS:** Resources, Writing – Review & Editing **AD:** Resources, Writing – Review & Editing **RP:** Resources, Data Curation, Writing – Review & Editing **SP:** Resources, Writing – Review & Editing **SA:** Resources, Writing – Review & Editing **IE:** Resources, Writing – Review & Editing **RR:** Resources, Writing – Review & Editing **RL:** Resources, Validation, Data Curation **AV:** Resources, Validation, Data Curation **MV:** Resources, Validation, Data Curation **MSC:** Resources, Writing – Review & Editing **DF:** Resources, Validation, Formal analysis, Investigation, Data Curation, Writing – Review & Editing **CMP:** Conceptualization, Methodology, Validation, Formal analysis, Investigation, Data Curation, Writing – Review & Editing, Visualization **AV:** Conceptualization, Methodology, Validation, Formal analysis, Investigation, Data Curation, Writing – Review & Editing, Visualization **XS:** Conceptualization, Software, Formal analysis, Data Curation, Writing – Review & Editing, Visualization **EN:** Conceptualization, Methodology, Validation, Formal analysis, Investigation, Resources, Data Curation, Writing – Original Draft, Writing – Review & Editing, Visualization, Supervision, Project administration, Funding acquisition.

## References

1. Delgermaa V, Takahashi K, Park EK, L. GV, Hara T, Sorahan T. Global mesothelioma deaths reported to the World Health Organization between 1994 and 2008. Bull World Health Organ 2011;89:716–24, 24A-24C.

2. Chen T, Sun XM, Wu L. High Time for Complete Ban on Asbestos Use in Developing Countries. JAMA oncology 2019;5:779–80.

3. Woolhouse I, Bishop L, Darlison L, de Fonseka D, Edey A, Edwards J, et al. BTS guideline for the investigation and management of malignant pleural mesothelioma. BMJ Open Respir Res 2018;5:e000266.

4. Vogelzang NJ, Rusthoven JJ, Symanowski J, Denham C, Kaukel E, Ruffie P, et al. Phase III study of pemetrexed in combination with cisplatin versus cisplatin alone in patients with malignant pleural mesothelioma. J Clin Oncol 2003;21:2636–44.

5. Zalcman G, Mazieres J, Margery J, Greillier L, Audigier-Valette C, Moro-Sibilot D, et al. Bevacizumab for newly diagnosed pleural mesothelioma in the Mesothelioma Avastin Cisplatin Pemetrexed Study (MAPS): a randomised, controlled, open-label, phase 3 trial. Lancet 2016;387:1405–14.

6. Popat S, Curioni-Fontecedro A, Polydoropoulou V, Shah R, O’+Brien M, Pope A, et al. A multicentre randomized phase III trial comparing pembrolizumab (P) vs single agent chemotherapy (CT) for advanced pre-treated malignant pleural mesothelioma (MPM): Results from the European Thoracic Oncology Platform (ETOP 9-15) PROMISE-meso trial. Ann Oncol 2019;30:v931. Abstract 1665.

7. Baas P, Scherpereel A, Nowak A, Fujimoto N, Peters S, Tsao A, et al. First-line nivolumab + ipilimumab vs chemotherapy in unresectable malignant pleural mesothelioma: CHECKMATE 743. Presented at WCLC 2020 Virtual Presidential Symposium on 08 August 2020.

8. Hmeljak J, Sanchez-Vega F, Hoadley KA, Shih J, Stewart C, Heiman D, et al. Integrative Molecular Characterization of Malignant Pleural Mesothelioma. Cancer Discov 2018;8:1548–65.

9. Lopez-Rios F, Chuai S, Flores R, Shimizu S, Ohno T, Wakahara K, et al. Global gene expression profiling of pleural mesotheliomas: overexpression of aurora kinases and P16/CDKN2A deletion as prognostic factors and critical evaluation of microarray-based prognostic prediction. Cancer Res 2006;66:2970–9.

10. Dacic S, Kothmaier H, Land S, Shuai Y, Halbwedl I, Morbini P, et al. Prognostic significance of p16/cdkn2a loss in pleural malignant mesotheliomas. Virchows Arch 2008;453:627–35.

11. Hylebos M, Van Camp G, van Meerbeeck JP, Op de Beeck K. The Genetic Landscape of Malignant Pleural Mesothelioma: Results from Massively Parallel Sequencing. Journal of thoracic oncology: official publication of the International Association for the Study of Lung Cancer 2016;11:1615–26.

12. Frizelle SP, Grim J, Zhou J, Gupta P, Curiel DT, Geradts J, et al. Re-expression of p16INK4a in mesothelioma cells results in cell cycle arrest, cell death, tumor suppression and tumor regression. Oncogene 1998;16:3087–95.

13. Middleton G, Fletcher P, Popat S, Savage J, Summers Y, Greystoke A, et al. The National Lung Matrix Trial of personalized therapy in lung cancer. Nature 2020;583:807–12.

14. Sherr CJ, Beach D, Shapiro GI. Targeting CDK4 and CDK6: From Discovery to Therapy. Cancer Discov 2016;6:353–67.

15. Finn RS, Crown JP, Lang I, Boer K, Bondarenko IM, Kulyk SO, et al. The cyclin-dependent kinase 4/6 inhibitor palbociclib in combination with letrozole versus letrozole alone as first-line treatment of oestrogen receptor-positive, HER2-negative, advanced breast cancer (PALOMA-1/TRIO-18): a randomised phase 2 study. The Lancet Oncology 2015;16:25–35.

16. Turner NC, Ro J, Andre F, Loi S, Verma S, Iwata H, et al. Palbociclib in Hormone-Receptor-Positive Advanced Breast Cancer. N Engl J Med 2015;373:209–19.

17. Bonelli MA, Digiacomo G, Fumarola C, Alfieri R, Quaini F, Falco A, et al. Combined Inhibition of CDK4/6 and PI3K/AKT/mTOR Pathways Induces a Synergistic Anti-Tumor Effect in Malignant Pleural Mesothelioma Cells. Neoplasia 2017;19:637–48.

18. Campisi J, d’+Adda di Fagagna F. Cellular senescence: when bad things happen to good cells. Nat Rev Mol Cell Biol 2007;8:729–40.

19. Finn RS, Dering J, Conklin D, Kalous O, Cohen DJ, Desai AJ, et al. PD 0332991, a selective cyclin D kinase 4/6 inhibitor, preferentially inhibits proliferation of luminal estrogen receptor-positive human breast cancer cell lines in vitro. Breast Cancer Res 2009;11:R77.

20. Goel S, DeCristo MJ, Watt AC, BrinJones H, Sceneay J, Li BB, et al. CDK4/6 inhibition triggers anti-tumour immunity. Nature 2017;548:471–5.

21. Salvador-Barbero B, Alvarez-Fernandez M, Zapatero-Solana E, El Bakkali A, Menendez MDC, Lopez-Casas PP, et al. CDK4/6 Inhibitors Impair Recovery from Cytotoxic Chemotherapy in Pancreatic Adenocarcinoma. Cancer Cell 2020;37:340–53 e6.

22. Ruscetti M, Leibold J, Bott MJ, Fennell M, Kulick A, Salgado NR, et al. NK cell-mediated cytotoxicity contributes to tumor control by a cytostatic drug combination. Science 2018;362:1416–22.

23. Wagner V, Gil J. Senescence as a therapeutically relevant response to CDK4/6 inhibitors. Oncogene 2020;39:5165–76.

24. Bezzi M, Teo SX, Muller J, Mok WC, Sahu SK, Vardy LA, et al. Regulation of constitutive and alternative splicing by PRMT5 reveals a role for Mdm4 pre-mRNA in sensing defects in the spliceosomal machinery. Genes Dev 2013;27:1903–16.

25. AbuHammad S, Cullinane C, Martin C, Bacolas Z, Ward T, Chen H, et al. Regulation of PRMT5-MDM4 axis is critical in the response to CDK4/6 inhibitors in melanoma. Proc Natl Acad Sci U S A 2019;116:17990–8000.

26. Varin E, Denoyelle C, Brotin E, Meryet-Figuiere M, Giffard F, Abeilard E, et al. Downregulation of Bcl-xL and Mcl-1 is sufficient to induce cell death in mesothelioma cells highly refractory to conventional chemotherapy. Carcinogenesis 2010;31:984–93.

27. Oie HK, Russell EK, Carney DN, Gazdar AF. Cell culture methods for the establishment of the NCI series of lung cancer cell lines. J Cell Biochem Suppl 1996;24:24-31.28.

28. Nadal E, Chen G, Gallegos M, Lin L, Ferrer-Torres D, Truini A, et al. Epigenetic inactivation of microRNA-34b/c predicts poor disease-free survival in early-stage lung adenocarcinoma. Clin Cancer Res 2013;19:6842–52.

29. Bollard J, Miguela V, Ruiz de Galarreta M, Venkatesh A, Bian CB, Roberto MP, et al. Palbociclib (PD-0332991), a selective CDK4/6 inhibitor, restricts tumour growth in preclinical models of hepatocellular carcinoma. Gut 2017;66:1286–96.

30. Puschel F, Munoz-Pinedo C. Measuring the Activation of Cell Death Pathways upon Inhibition of Metabolism. Methods Mol Biol 2019;1862:163–72.

31. Core Team. R: A language and environment for statistical computing. R Foundation for Statistical Computing, Vienna, Austria URL: https://wwwR-projectorg/ 2017.

32. Ambrogio C, Carmona FJ, Vidal A, Falcone M, Nieto P, Romero OA, et al. Modeling lung cancer evolution and preclinical response by orthotopic mouse allografts. Cancer Res 2014;74:5978–88.

33. Bueno R, Stawiski EW, Goldstein LD, Durinck S, De Rienzo A, Modrusan Z, et al. Comprehensive genomic analysis of malignant pleural mesothelioma identifies recurrent mutations, gene fusions and splicing alterations. Nature genetics 2016;48:407–16.

