## supplementary figures for "Efficacy of CDK4/6 inhibitors in preclinical models of malignant pleural mesothelioma"

#### Suppl. Figure S1

A

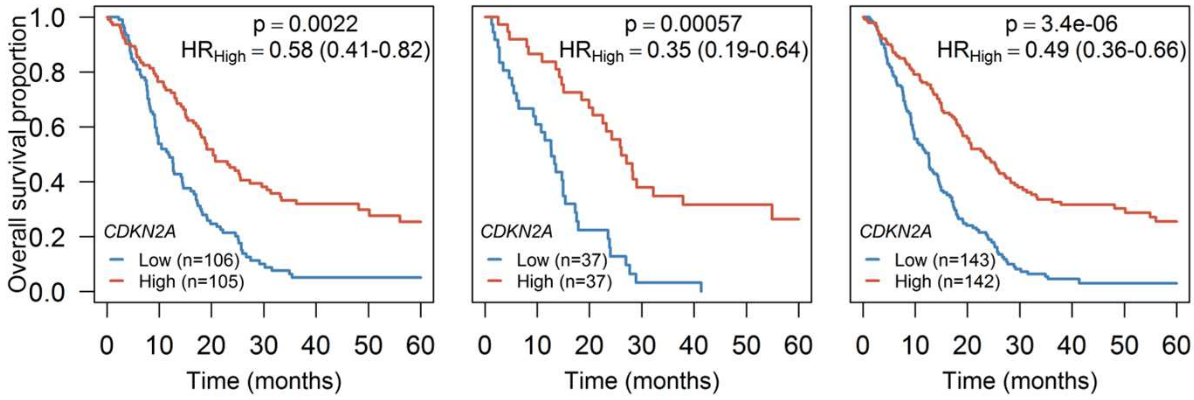


B

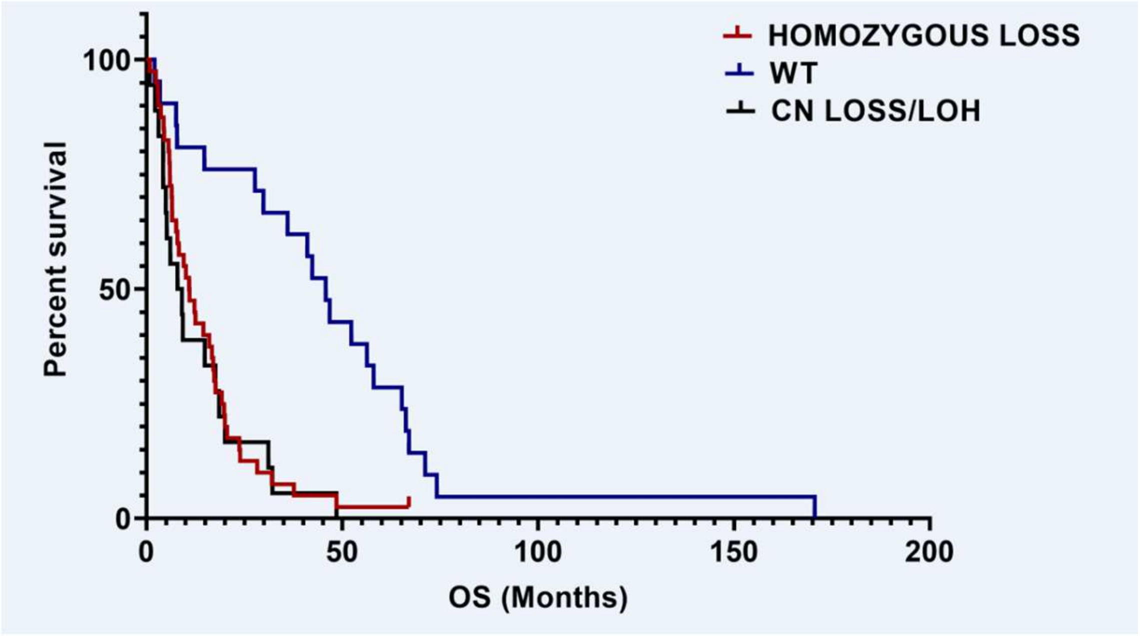

### Suppl. Figure S2

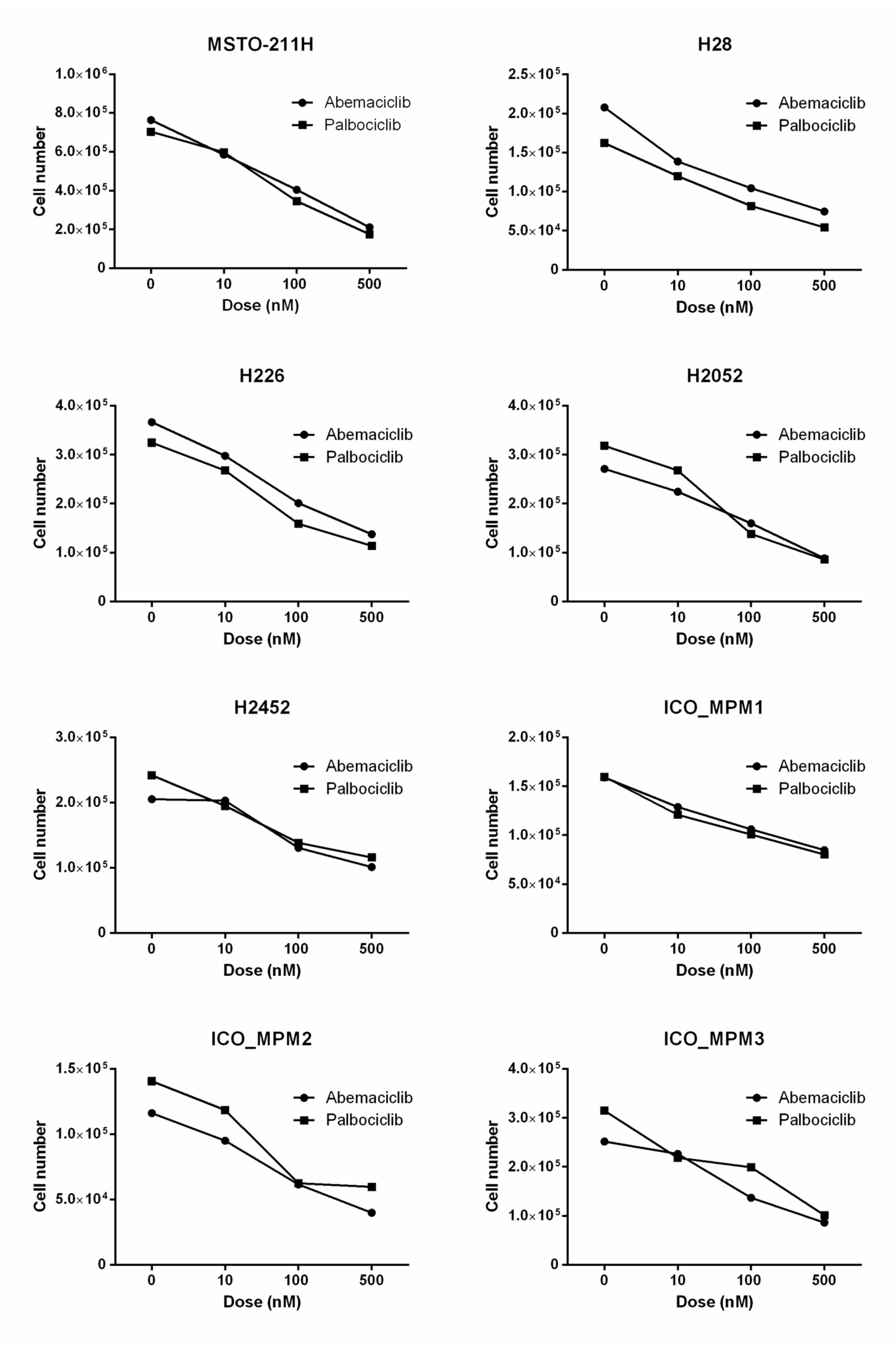

### Suppl. Figure S3

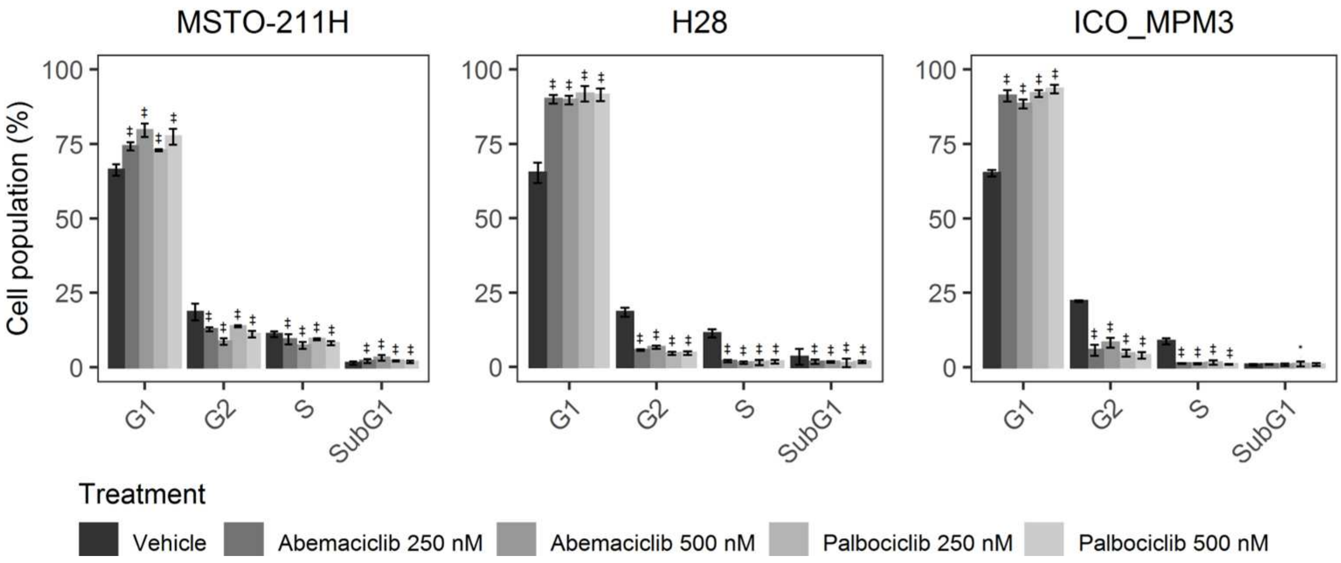

### Suppl. Figure S4

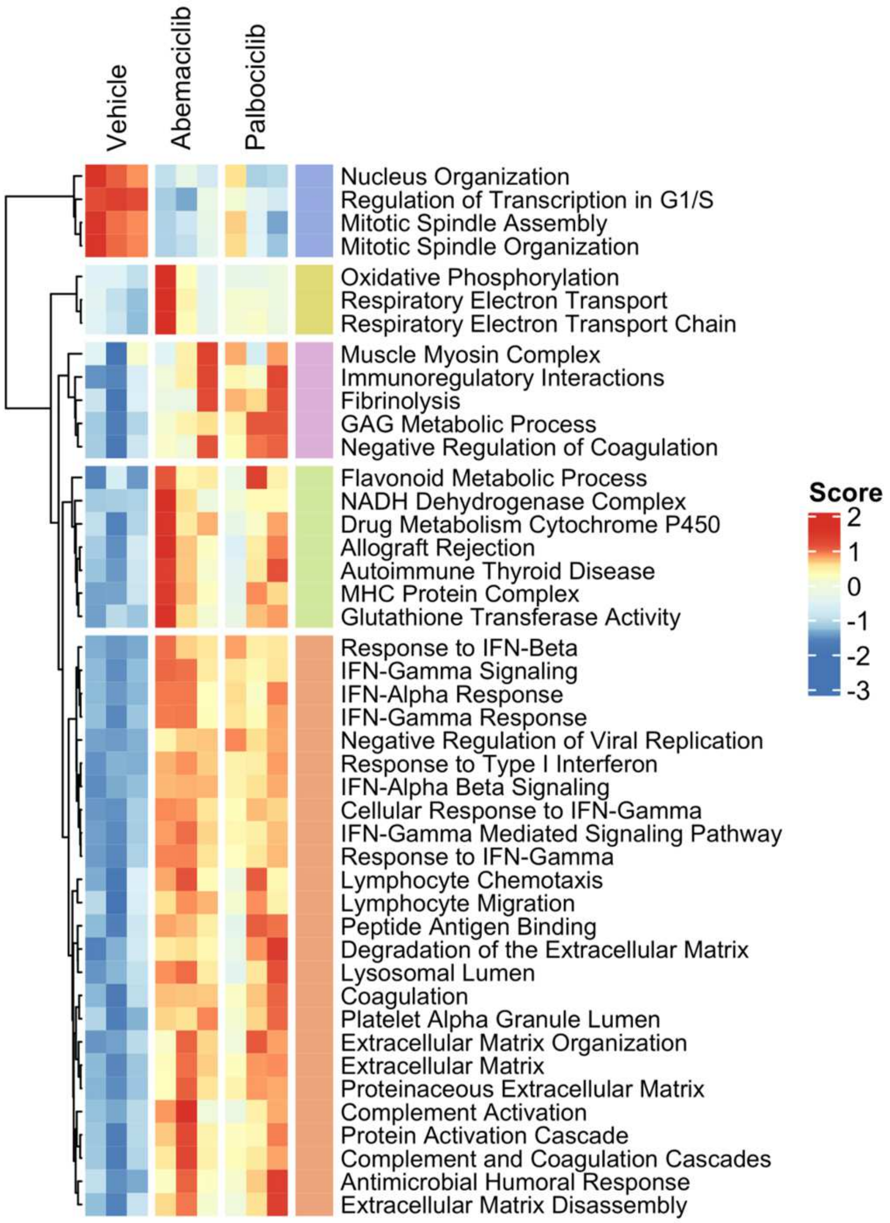

### Suppl. Figure S5

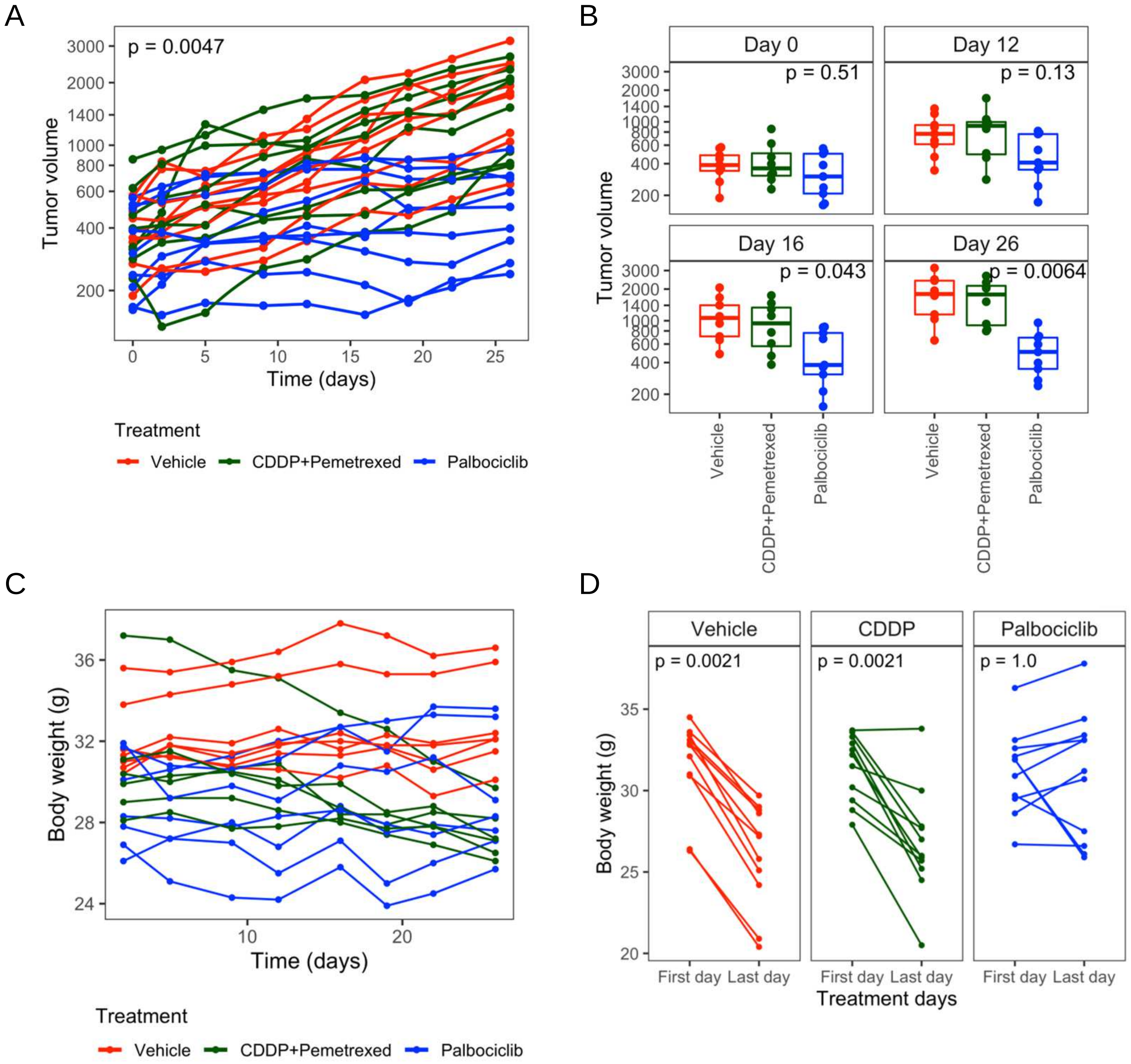

**Suppl. Figure S6**

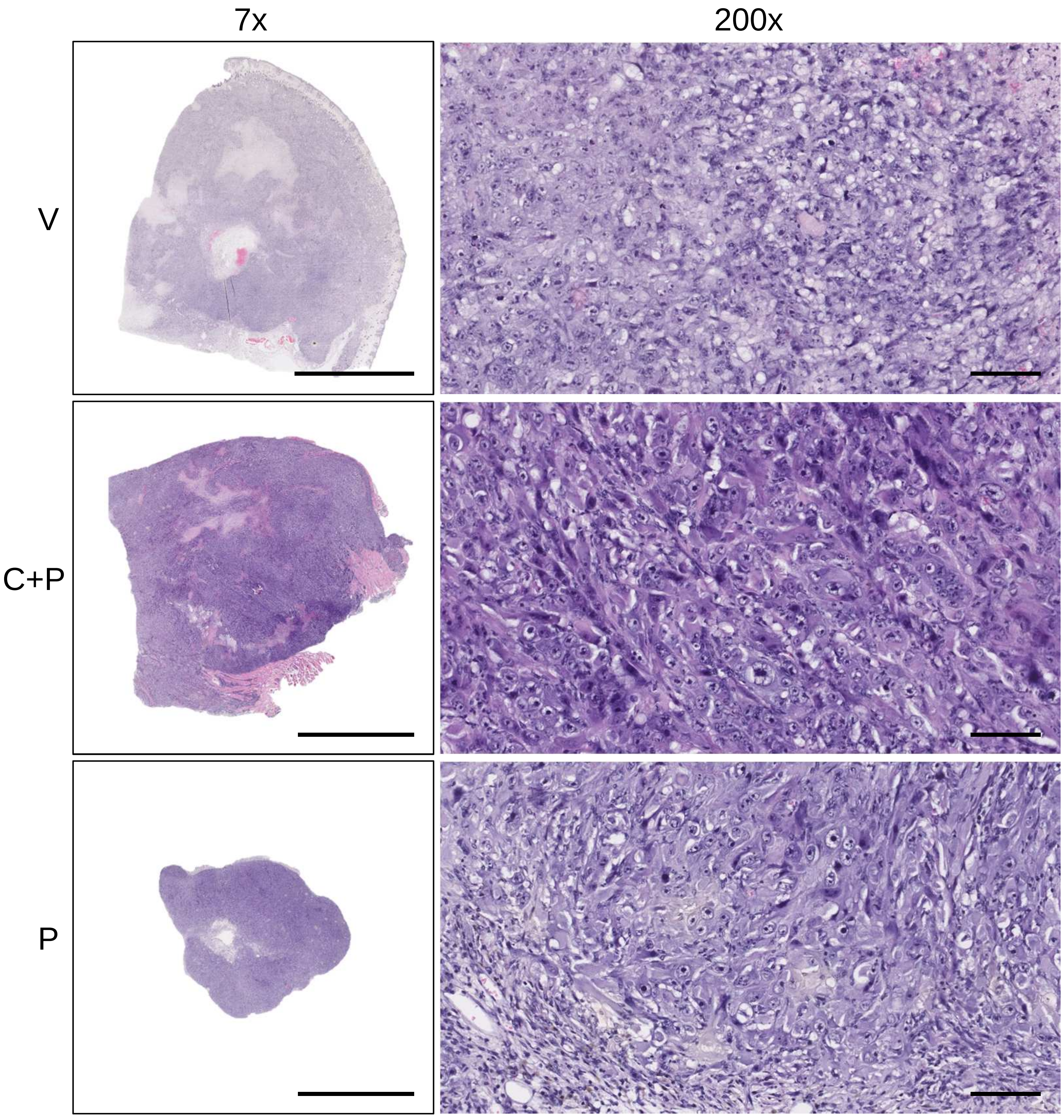

### Suppl. Figure S7

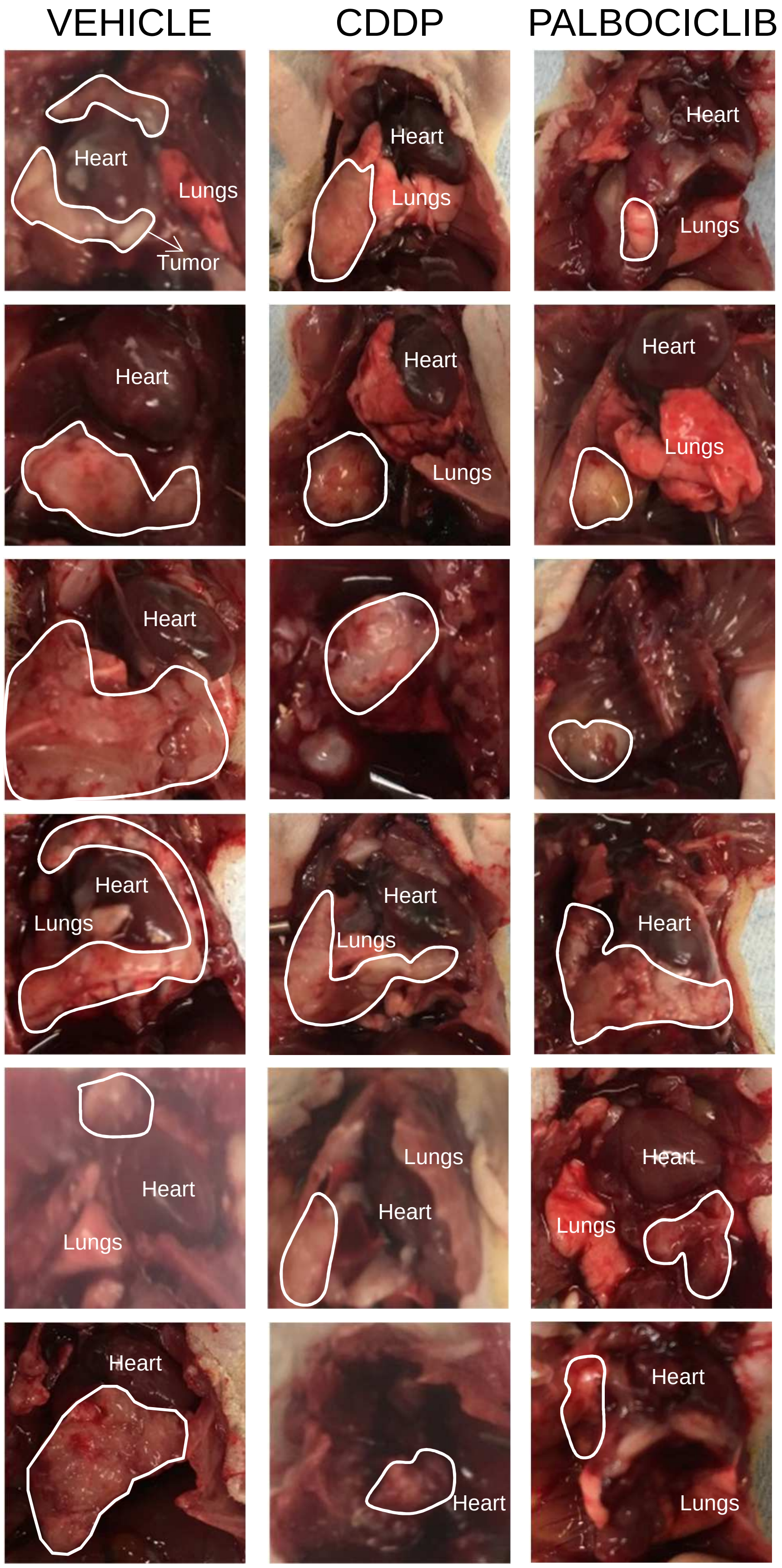

**Suppl. Figure S8**

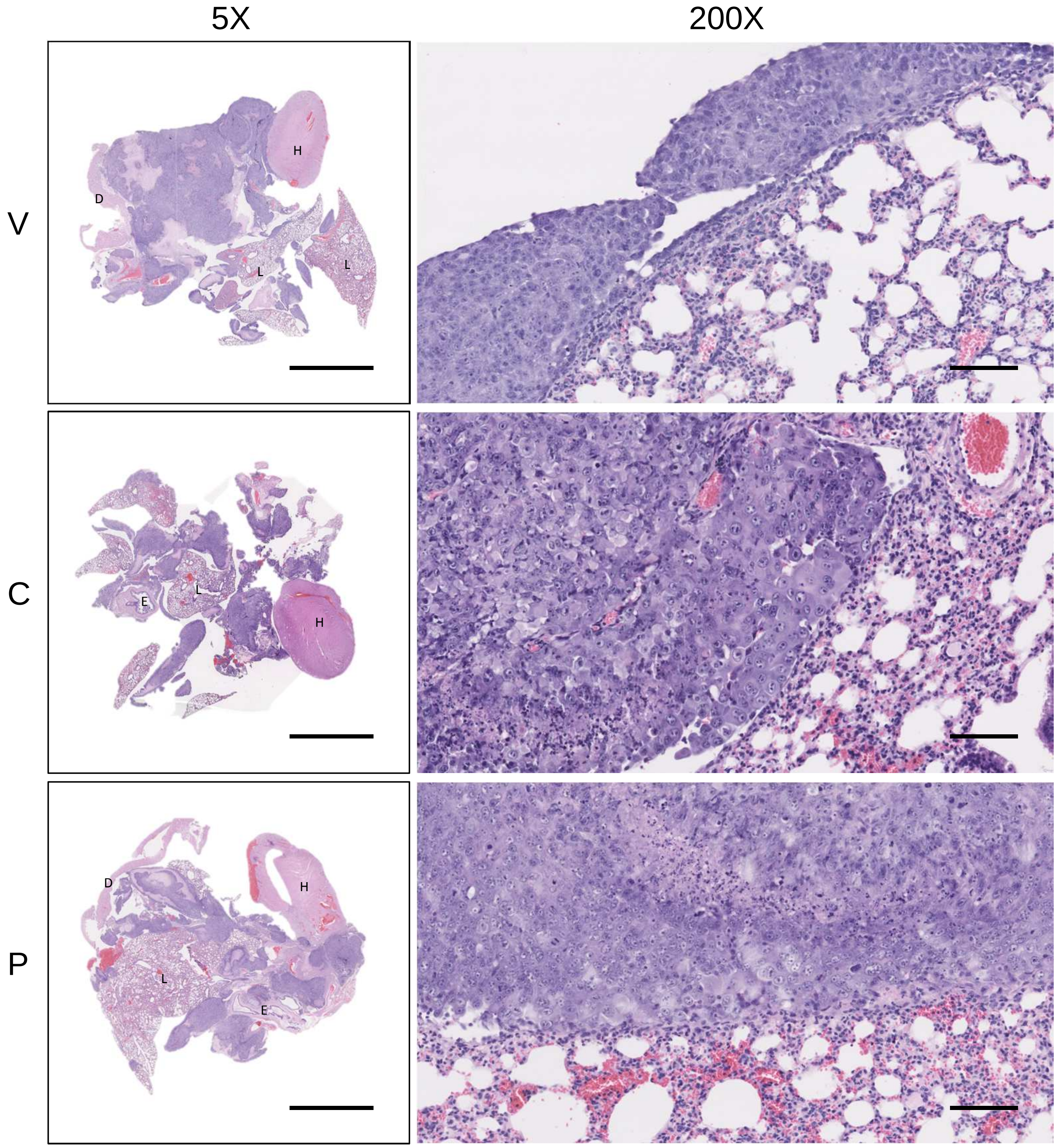
