## supplementary figure legends for "Efficacy of CDK4/6 inhibitors in preclinical models of malignant pleural mesothelioma"

**Suppl. Figure S1.** **Effects of *CDKN2A* expression and copy number alterations on overall survival in MPM patients** A) Kaplan-Meier plots of overall survival (OS) in MPM patients according to *CDKN2A* gene expression levels based on data obtained from Bueno et al. (left column) and Hmeljak et al. (middle column) cohorts or the combination of both (right column). Low levels of *CDKN2A* (blue line) were significantly associated with poor OS in patients with MPM. In each cohort, the high and low expression levels were defined based upon the median. P-values and hazard ratios (HR) were calculated by likelihood ratio test and multivariate Cox regression analysis respectively. B) Kaplan Meier curve for overall survival in surgical samples (n=79) in patients with MPM according to the *CDKN2A* copy number status.

**Suppl. Figure S2. Drug response curves for both CDK4/6 inhibitors in 5 commercial (MSTO, 211H, H28, H226, H2052 and H2452) and 3 primary patient-dervied (ICO_MPM1, ICO_MPM2 and ICO_MPM3) MPM cell lines.** Response to treatment with varying concentrations (0, 10, 100 and 500 nM) of abemaciclib or palbociclib for 72 hours.

**Suppl. Figure S3.** **Quantitative analysis of the cell cycle populations after treatment with vehicle (medium) and 250 and 500 nM of abemaciclib or palbociclib for 24 hours.** In MSTO-211H, H28 and ICO_MPM3, the percentage of cell population in G1-phase significantly increased, while the percentage of G2 and S cell cycle populations significantly decreased respect to control cells. An irrelevant 1% to 3% variation was observed in the percentage of subG1 population indicating that drug treatment was not associated with apoptosis induction. Cell cycle experiments were analyzed using proportion tests based on all the cells counted in the abovementioned experiments. P-values were adjusted using FDR.

**Suppl. Figure S4. Heatmap of single-sample GSEA scores from significant pathways in pre-ranked GSEA (time-course).** Heatmap showing enrichment score of pathways up and down-regulated in MSTO-211H control cells (left columns) and compared with cells treated with either abemaciclib 250 nM (middle columns) or palbociclib 250 nM (right columns). Significant pathways were defined as those with FWER < 0.05 in pre-ranked GSEAs comparing each treatment to the control using RNA sequencing data at 72 hours. (GSEA: gene set enrichment analysis; FWER: family-wise error rate).

**Suppl. Figure S5**. **Effects of treatments in tumor volume and body weight.** (A) Detailed plot showing the tumor volume variation from each mouse along the in vivo subcutaneous xenograft tumor growth experiment. (B) Differences in tumor volume for each day were compared using Kruskal-Wallis test and adjusted by FDR and were significantly different at day 16. Summary of the body weight values among first and last day of treatment from all mice in in vivo (C) subcutaneous and (D) orthotopic tumor xenograft growth experiments. Palbociclib did not exert any substantial change in the mice body weight, and it did not either in in vivo xenograft tumor growth experiment. Differences were evaluated by Mann-Whitney test and adjusted for FDR.

**Supplementary figure S6.** **Histologic appearance of the subcutaneous tumors.** H/E images of subcutaneous tumors after treatment with vehicle (V), CDDP + Pemetrexed (C+P) and Palbociclib (P). Original magnification of the left pictures: 7x, scale bars: 5 mm; original magnification of the right pictures: 200x, scale bars: 100 µm.

**Suppl. Figure S7.** **Macroscopic images of the orthotopic tumors in the mouse thoracic cavity after treatments.** The orthotopic xenograft mouse model was stablished by implanting MSTO-211H tumors, previously grown subcutaneously in athymic nude mice treated with chemotherapy, in the pleural cavity of athymic mice. Representative images of the different treatments (6 for each treatment) in the orthotopic experiment showed the growth of tumors in the pleural cavity. The area delimited with a white line denote tumor mass.

**Suppl. Figure S8.** **Histologic appearance of the orthotopic tumors.** Tumor cells orthotopically implanted in the thoracic cavity are growing on the pleural, pericardial or adventitial surfaces of the lungs, heart or esophagus reproducing the growth pattern of malignant mesothelioma. Left pictures show a panoramic view of the thoracic structures (H&E, 5x, scale bars: 5 mm); right pictures show a detail of the tumor growing on the visceral pleura (H&E, 200x, scale bars: 100 µm). Figure legends: C: cisplatin; D: diaphragm; E: esophagus, H: heart, L: lung; P: palbociclib and V: vehicle.
