## supplementary methods for "Efficacy of CDK4/6 inhibitors in preclinical models of malignant pleural mesothelioma"

**Cell culture and cell lines**

H28, H2452, H2052, MSTO-211H and H226 cells were purchased from the American Type Culture Collection (ATCC, Manassas, Virginia) and cultured in Roswell Park Memorial Institute 1640 medium (RPMI-1640, Life Technologies, Paisley, Great Britain) supplemented with 10% fetal bovine serum (FBS, Life Technologies) and penicillin-streptomycin (100 U/ml, Life Technologies). ICO_MPM1, ICO_MPM2 and ICO_MPM3 are primary cell lines derived from pleural effusions of three patients diagnosed with malignant pleural mesothelioma (MPM) at the Hospital Universitari de Bellvitge – Catalan Institute of Oncology (L’Hospitalet, Barcelona, Spain). Samples were obtained after patients signed the informed consent approved by the local Ethical Committee (PR152/14). Pleural effusions were collected by pleural tap into sterile bottles and tumor cells were isolated and cultured as previously described [1]. Briefly, cells were spun down in sterile falcon 50 ml centrifuge tubes at 1,250 rpm for 15 minutes. Pelleted cells were then seeded into 150 mm^2^ petri dishes in 20 ml of basal medium RPMI-1640 supplemented with 5% FBS, 100 U/ml penicillin-streptomycin, 2 mM glutamine (Life Technologies), 10 mM Hepes (Sigma-Aldrich, St. Louis, Missouri), 20 µg/ml insulin (Sigma-Aldrich), 10 µg/ml transferrin (Sigma-Aldrich), 25 nM sodium selenite (Sigma-Aldrich), 50 nM hydrocortisone (Sigma-Aldrich), 1 ng/ml epidermal growth factor (EGF, Bionova Cientifica, Rocky Hill, New Jersey), 10 µM ethanolamine (Sigma-Aldrich), 10 µM phosphorylethanolamine (Sigma-Aldrich), 20 µg/ml triiodothyronine (Sigma-Aldrich) and 0.5 mM sodium pyruvate (Sigma-Aldrich). Tumor cells were isolated from other culture cell populations by selective trypsinization along passages. All cells were incubated and maintained at 37˚C in humidified chamber containing 5% CO_2_.

**Western blot analysis**

MPM commercial and primary cell lines were lysed using RIPA buffer (Thermo Fisher Scientific, Waltham, Massachusetts) containing inhibitors of proteases and phosphatases (Complete and PhosSTOP, Roche, Basel, Switzerland). Protein concentrations were measured using BCA protein assay kit (Thermo Scientific) and 10 μg protein was loaded onto pre-cast 4-20% Mini-Protean TGX stain free gels (Bio-Rad, Hemel Hempstead, Great Britain) for electrophoresis and then transferred onto nitrocellulose membranes (Bio-Rad). Membranes were blocked with 5% non-fat milk (Nestle, Vevey, Switzerland) in tris-buffered saline (TBS) containing 0.1% Tween® 20 (TBS-T) at pH 7.4, for 1 hour at room temperature (RT). Primary antibodies (1/1000) were incubated overnight at 4˚C in TBS-T and 1% non-fat milk. After 3 washes with TBS-T, polyclonal goat anti-rabbit horseradish peroxidase-conjugated secondary antibody (1/8000; Sigma-Aldrich) was incubated at RT for 1 hour. Immunoreactive bands were detected using ECL SelectTM Western Blotting Detection Reagent (GE Healthcare, Chicago, Illinois) and digitized using the ChemiDoc^TM^ Touch Imaging System (Bio-Rad).

**Cell viability assays (cell counting and colony formation assays)**

For cell counting studies, cells were plated into 6-well plates and treated at 0, 10, 100 and 500 nM of abemaciclib or palbociclib for 72 hours. Cells were then collected and counted using a Neubauer chamber (VWR International, Radnor, Pennsylvania). Trypan blue was used as a stain to quantify live cells by labeling dead cells exclusively. A minimum of three replicates were performed for each condition. The drugs were added when cells were attached to the plate and medium was not changed for the length of the treatment.

For colony formation assay, 1000 cells were seeded into 6-well plates and treated with 0, 250 and 500 nM of abemaciclib or palbociclib for 15 days. Then, cells were washed with phosphate buffer saline (PBS), fixed with methanol for 15 minutes and stained with 0.1% crystal violet for 20 minutes. Finally, plates were washed under tap water and allowed to dry overnight prior to being scanned. Each assay was repeated at least three times. Both cell viability assays were assessed using Wilcoxon signed rank tests comparing each treatment with vehicle.

**Cell cycle and apoptosis analysis**

For cell cycle evaluation, 1 x 10^5^ cells per well were plated in 6-well plates and treated at 0, 250 and 500 nM of abemaciclib or palbociclib for 24 hours. Then, cells were incubated in cell lysis solution with trypsin. After 10 minutes, trypsin was inactivated with trypsin-inactivating solution and lysed cells were stained with propidium iodide (PI; 40 µg/ml; Sigma) for 20 minutes at RT and protected from light. Finally, DNA content was analyzed by FACS. A minimum of 10,000 events were analyzed on the Beckman Coulter Gallios Flow cytometer (Beckman Coulter Inc, Brea, California) and data were analyzed with FlowJo v10.0 (FlowJo LLC, Ashland, Oregon). A minimum of three biological replicates were performed. Statistical significance of the cell cycle experiments was analyzed using proportion tests based on all the cells counted in the abovementioned experiments. P-values were adjusted using false discovery rate (FDR).

For cell death evaluation, 1 x 10^5^ cells/well were seeded in 6-well plates and treated with 0, 250 and 500 nM abemaciclib or palbociclib for 72 hours. Then, cells were subjected to trypsinization, washed with PBS, pelleted, stained with propidium iodide (PI) at 0.5µg/ml in PBS EDTA (1mM) and incubated for 20 minutes prior to FACS analysis. Cell death was measured by using PI incorporation. In order to quantify non-apoptotic cell death, cells were treated with QVD (10 µM; ApexBio, Houston, Texas) in DMSO and the measured cell death values were compared to those treated with a vehicle control (total cell death); the difference was presented as the rate of apoptotic cell death. Statistical differences among treatment and vehicle conditions in cell death experiment were evaluated using Mann-Whitney U test for each comparison and Kruskal-Wallis test if 3 conditions were simultaneously tested and adjusted afterwards using FDR.

**Measurement of cellular senescence**

Senescence was measured by using the SA-β-gal staining kit (Cell Signaling Technology), following the manufacturer’s protocol. Briefly, cells were seeded in 6-well plates (1 x 10^5^ cells/well) and incubated with 0, 250 and 500 nM abemaciclib or palbociclib for 72 hours. After washing with PBS twice, fixative solution was added for 15 minutes at RT. Plates were then washed again and the β-Galactosidase staining solution was added. After overnight incubation at 37˚C without CO_2_, the number of SA-β-gal positive cells (blue stained) was evaluated by cell counting in five randomly chosen microscope fields (200 magnification). Statistical significance of the cellular senescence experiments was evaluated using proportion tests based on all the cells counted in the abovementioned experiments. P-values were adjusted using FDR.

***In vivo* subcutaneous xenograft tumor growth experiments**

The following experiments were performed under the 9111 experimental procedure and in accordance with protocols approved by Animal Research Ethics Committee of the Bellvitge Biomedical Research Institute Animal Facility (Registration number B9900010). Four x 10^6^ MSTO-211H cells growing exponentially were suspended in 300 $\mu$l PBS and subcutaneously inoculated into the right flanks of 30 four-week-old male athymic nude mice (Envigo, Indiana, Indianapolis). Once the tumors reached a homogeneous average volume size of 300–400 mm^3^, mice (n=28) were randomly assigned into four groups (n = 7 mice/per group) and treated with: i) vehicle, 200 μl of 0.05 N sodium lactate pH 4.0 by oral gavage 5 days over 7 days; ii) combination of cisplatin, 3.5 mg/kg administered intraperitoneally once a week plus pemetrexed, 100 mg/kg intraperitoneally twice a week and iii) single palbociclib, 150 mg/kg by oral gavage 5 days over 7 days. To evaluate efficacy, tumor volumes (V =π/6 x L x W^2^) were measured twice per week with calipers and the weight of each animal was measured every day. After 26 days of treatment, mice were euthanized by cervical dislocation and the tumors were excised, weighed and processed for histologic and for RNA studies following standard protocols. The mean volume + SD were calculated. Using R software v.3.5.0 [2], a linear mixed effects model was fitted with a random intercept for each mouse and within-subject correlation as autoregressive of order one to assess the significance of treatment independently of day in explaining variations in tumor growth. A longitudinal analysis testing for differences in treatment slopes was done using analysis of covariance. Tumor volume differences among treatments for each day were assessed using Kruskal-Wallis tests, with FDR adjustment.

***In vivo* orthotopic xenograft tumor growth experiments**

Tumors grown subcutaneously in athymic nude mice (Charles River Laboratories Inc) treated and with cisplatin plus pemetrexed were aseptically isolated the same day of sacrifice, minced in small tumor fragments and kept in DMEM supplemented with 10% FBS (Life Technologies) plus 50 U/mL penicillin and 50 mg/mL streptomycin. Small solid fragments of the tumors were implanted orthotopically in the lung of 35 athymic nude mice following our previously reported procedures [3]. Briefly, mice were anesthetized with a continuous flow of 1% to 3% isoflurane/oxygen mixture (2 L/min) and subjected to right thoracotomy. Mice were situated in left lateral decubitus position, and a small transverse skin incision (5–8 mm) was made in the right chest wall. Chest muscles were separated by a sharp dissection and costal and intercostal muscles were exposed. An intercostal incision of 2 to 4 mm on the third or fourth rib on the chest wall was made and a small tumor piece of 2 to 4 mm^3^ was introduced into the chest cavity and the tumor specimens were anchored to the lung surface with Prolene 7.0 suture. Next, the chest wall incision was closed with surgery staples, and finally chest muscles and skin were closed. The waiting time between tumor implantation and the beginning of treatments was of 2 weeks based on our previous orthotopic xenograft MPM model experience. Thirty-three alive mice were randomized into three groups (n = 11 mice/group) and treated with: i) vehicle, 200 μl of 0.05 N sodium lactate pH 4.0 by oral gavage 5 days over 7 days; ii) cisplatin, 3.5 mg/kg intraperitoneally once a week; or iii) palbociclib, 150 mg/Kg by oral gavage 5 days over 7 days during forty days. Mice were weighed daily and monitored for the presence of breathing problems. Beyond 40 days, all live mice remained untreated until human endpoint. Orthotopic tumors were collected from euthanized mice when presenting breathing problems and processed for histological studies following standard protocols. Kaplan-Meier curves for each cohort of mice were calculated and differences between groups were compared using Cox proportional hazards model. Differences in body weight were analyzed using Mann-Whitney U test comparing first and last day of treatment in each treatment and adjusted with FDR.

***In silico* analysis of publicly available RNA-sequencing data**

Public data from RNA-seq cohorts published by Bueno et al. [4] and Hmeljak et al. [5] was obtained from European Genome-Phenome Archive [6] and TCGA2BED [7] repositories respectively. For the Bueno et al. data, raw reads were pre-processed using Trimmomatic v.0.32 [8] to perform quality trimming and adapters’ removal, and an in-house Python script was used to remove reads with undetermined bases. Subsequently, reads were aligned to GENCODE v26 genome (GRCh38) using STAR v.2.5.3.a [9] and quantified with RSEM v.1.3.0 [10] to obtain transcripts per million (TPM). For Hmeljak et al. data, data was downloaded already in TPM from TCGA2BED repository.

Log-transformed TPMs (Log_2_(TPM)) data for both cohorts was used in the analysis. Gene expression status was stratified using the median, and Cox proportional-hazards models adjusted for sex, stage, age and histology were fitted to assess differences in survival using R software [2].

**RNA extraction and quality assessment**

Total RNA from cells and xenografted tumors in mice was extracted by TRIzol (Life Technologies) and isolated using the RNeasy Mini Kit (Qiagen, Hilden, Germany) respectively, following the manufacturer’s protocol. Genomic DNA digestion was performed using RNase-Free DNase Set (Qiagen) during RNA purification. RNA quantity and purity were measured using NanoDrop1 ND-1000 spectrophotometer (Thermo Scientific, UK). All A260/A280 and A260/A230 absorbance ratios were between 1.8–2.0 and 2.0–2.2, respectively. RNA quality and integrity were verified using the Eukaryote Total RNA Nano 6000 LabChip assay (Agilent Technologies, Santa Clara, California) on the Agilent 2100 Bioanalyzer. All RNA samples showed an RNA Integrity Number (RIN) > 8.

**Whole Exome Sequencing (WES) analysis of patient-derived cell lines**

WES libraries were generated using the Nextera Rapid Capture Enrichment library preparation kit (Illumina, San Diego, California), according to the manufacturer’s recommendations and were sequenced on an Illumina HiSeq 2500 paired-end with 100 base pair long reads. WES data processing Sequencing reads were pre-processed with Trimmomatic v.0.38 [8] performing quality trimming and adapters’ removal. Moreover, removal of reads with undetermined bases was performed using an in-house Python script. Pre-processed reads were mapped to NCBI human genome GRCh38 using BWA-mem version 0.7.17. Somatic variant calling was done following Genome Analysis Toolkit (GATK) best practices [11]. Using GATK v.4.1.1 aligned reads were marked for duplicates; base recalibrated and finally variants were called using Mutect2 algorithm. The final set of variants, including only variants passing Mutect2 filters, in standard chromosomes, and null variant allele frequency in normal samples was annotated using wANNOVAR [12]. Driver variants were identified using Cancer Genome Interpreter [13].

**RNA sequencing (RNA-seq) analysis of patient-derived cell lines**

Total RNA was submitted to the Centro National de Análisis Genómico (CNAG, Barcelona, Spain), for RNA-Seq library preparation and sequencing. RNA-Seq libraries were prepared from the enriched RNA using the NEBNext Ultra Directional RNA library Prep kit (Illumina). Libraries were purified using AMPure XP beads (Beckman Coulter). Each library was quantified using Qubit (Thermo Fisher Scientific) and the size distribution assessed using the Agilent 2100 Bioanalyser (Agilent technologies). Libraries were pooled and assessed by a Qubit dsDNA HS assay kit and qPCR using the Illumina Library Quantification kit from Kapa (Roche) on a Roche Light Cycler ^®^480 Instrument II. Concentrations of template DNA were adjusted to 3 nM and denatured with 0.1 N NaOH. Denatured cDNA was further diluted to a final concentration of 300 pM. Clustering of the DNA templates was performed using HiSeq 3000/4000 paired end cluster Kit with a cBot cluster generation system (Illumina) according to the manufacturer’s instructions. Libraries were sequenced on an Illumina HiSeq 4000 using sequencing by synthesis chemistry to generate 2 x 150bp paired end reads. Raw RNA-seq reads were pre-processed using the same pipeline used for whole exome sequencing and then aligned to GENCODE v29 genome (GRCh38) using STAR v.2.6.0 [9]. Quantification of reads was performed with RSEM v.1.3.1 [10], and TPM were retrieved. All quality controls were done using FastQC version 0.11.7. Log_2_(TPM) were used to conduct a pre-ranked GSEA [14] to find associated pathways for each condition with respect to the control condition (vehicle-treated cells/mice). Included gene sets from MSigDB (v6.1) were KEGG, Biocarta, Reactome, Hallmarks, and Gene Ontology. Moreover, in order to evaluate the enrichment of significant gene sets (using a FWER threshold level of 0.05), a single-sample GSEA [15] was performed on all samples from the time course experiment. All statistical analyses were done using R software v.3.5.0 [2].

**References:**

1. Oie HK, Russell EK, Carney DN, Gazdar AF. Cell culture methods for the establishment of the NCI series of lung cancer cell  lines. *J Cell Biochem Suppl*. United States; **1996**;24:24–31.
2. Core Team (2017). R: A language and environment for statistical computing. R Foundation for Statistical Computing, Vienna, Austria. URL: <https://www.R-project.org/>.
3. Ambrogio C, Carmona FJ, Vidal A, Falcone M, Nieto P, Romero OA, *et al*. Modeling lung cancer evolution and preclinical response by orthotopic mouse allografts. *Cancer Res*. United States; **2014**;74:5978–88.
4. Bueno R, Stawiski EW, Goldstein LD, Durinck S, De Rienzo A, Modrusan Z, *et al*. Comprehensive genomic analysis of malignant pleural mesothelioma identifies recurrent mutations, gene fusions and splicing alterations. *Nat Genet*. United States; **2016**;48:407–16.
5. Hmeljak J, Sanchez-Vega F, Hoadley KA, Shih J, Stewart C, Heiman D, *et al*. Integrative molecular characterization of malignant pleural mesothelioma. *Cancer Discov*. **2018**;8:1548-65.
6. Lappalainen I, Almeida-King J, Kumanduri V, Senf A, Spalding JD, Ur-Rehman S, *et al*. The European Genome-phenome Archive of human data consented for biomedical research. *Nat Genet*. United States; **2015**;47:692-5.
7. Cumbo F, Fiscon G, Ceri S, Masseroli M, Weitschek E. TCGA2BED: extracting, extending, integrating, and querying The Cancer Genome Atlas. *BMC Bioinformatics*. England; **2017**;18:6.
8. Bolger AM, Lohse M, Usadel B. Trimmomatic: a flexible trimmer for Illumina sequence data. *Bioinformatics*. England; **2014**;30:2114–20.
9. Dobin A, Davis CA, Schlesinger F, Drenkow J, Zaleski C, Jha S, *et al*. STAR: ultrafast universal RNA-seq aligner. *Bioinformatics*. England; **2013**;29:15–21.
10. Li B, Dewey CN. RSEM: accurate transcript quantification from RNA-Seq data with or without a reference genome. *BMC Bioinformatics*. England; **2011**;12:323.
11. Van der Auwera GA, Carneiro MO, Hartl C, Poplin R, Del Angel G, Levy-Moonshine A, *et al*. From FastQ data to high confidence variant calls: the Genome Analysis Toolkit best practices pipeline. *Curr Protoc Bioinforma*. United States; **2013**;43:11.10.1-11.10.33.
12. Yang H, Wang K. Genomic variant annotation and prioritization with ANNOVAR and wANNOVAR. *Nat Protoc*. England; **2015**;10:1556–66.
13. Tamborero D, Rubio-Perez C, Deu-Pons J, Schroeder MP, Vivancos A, Rovira A, *et al*. Cancer Genome Interpreter annotates the biological and clinical relevance of tumor  alterations. *Genome Med*. **2018**;10:25.
14. Subramanian A, Tamayo P, Mootha VK, Mukherjee S, Ebert BL, Gillette MA, *et al*. Gene set enrichment analysis: a knowledge-based approach for interpreting genome-wide expression profiles. *Proc Natl Acad Sci U S A*. United States; **2005**;102:15545–50.
15. Hanzelmann S, Castelo R, Guinney J. GSVA: gene set variation analysis for microarray and RNA-seq data. *BMC Bioinformatics*. England; **2013**;14:7.
